# OncoBERT: Context-Aware Modeling of Somatic Mutations for Precision Oncology

**DOI:** 10.64898/2026.02.18.706658

**Authors:** Sushant Patkar, Noam Auslander, Stephanie Harmon, Peter Choyke, Baris Turkbey

## Abstract

Somatic mutation profiling is central to cancer diagnosis and treatment selection. However, most studies focus on individual actionable mutations, overlooking the broader mutational context that shapes tumor evolution and treatment response. Here, we introduce OncoBERT, a language model that learns contextual representations of somatic mutations from large-scale clinical sequencing data spanning >210,000 patients, 113 cancer types and 20 institutions. OncoBERT uncovers robust patient-specific mutational subtypes across diverse cohorts and targeted sequencing panels, revealing clinically meaningful mutation patterns that are associated with differential response to chemotherapy, targeted therapies, and immunotherapy. Importantly, integrating OncoBERT’s contextual representations with clinically approved biomarkers of immunotherapy response, such as tumor mutational burden (TMB) and microsatellite instability (MSI), significantly improved prediction of clinical benefit. By further incorporating matched tumor transcriptomic profiles, we linked OncoBERT-defined mutational subtypes to distinct cancer hallmark programs and tumor microenvironment states. Together, OncoBERT provides a scalable framework for deciphering somatic mutational landscapes, enabling improved patient stratification and advancing precision oncology.

## Introduction

Cancer is an evolutionary process, in part, driven by the accumulation of somatic mutations^1,2^. The advent of high-throughput sequencing technologies, such as whole-exome sequencing (WES) and targeted next-generation sequencing (e.g., MSK-IMPACT), has enabled large-scale profiling of somatic mutations across diverse tumor types^3–10^. These breakthroughs have ushered in a new generation of personalized cancer therapeutics focused on actionable mutations^11–18^. However, most cancer mutations do not occur in isolation^19^. Increasing evidence suggests that specific co-occurring mutations can significantly influence tumor aggressiveness and treatment response^20,21^. Therefore, a comprehensive understanding of how multiple cancer mutations interact, and influence patient outcomes is crucial for uncovering novel therapeutic targets and further advancing precision oncology. Several computational methods have been developed to study mutation patterns across cancer genomes^19,22–30^. These methods broadly fall under three categories: (i) pairwise co-mutation analysis methods (ii) network-based or pathway-based analysis methods and (iii) sequence-based analysis methods.

Pairwise co-mutation analyses focus on discovery of statistically significant co-mutated or mutually exclusive gene pairs that affect tumor fitness and patient survival. While simple and easy to interpret, these methods face notable limitations. First, the vast combinatorial space of potential mutation pairs makes exhaustive evaluation computationally expensive and susceptible to false positives, especially when considering relatively small dataset sizes^27,29,31^. Second, most pairwise analysis methods struggle with mutation data sparsity and are biased towards more frequently mutated oncogenes and tumor suppressors^22,32^. Third, these methods often fail to capture higher-order interactions involving three or more mutations, which may define clinically meaningful molecular subtypes^29^.

Network-based approaches alleviate some of these challenges by projecting mutation data onto predefined biological networks such as protein–protein interaction (PPI) networks^33–38^ to reveal frequently mutated cancer pathways. While effective, these strategies also have significant drawbacks. For instance, many network-based analysis approaches rely on non-negative matrix factorization (NMF), which does not scale well with large, heterogeneous datasets39,40. Furthermore, NMF’s parts-based decomposition approach often fails to effectively capture nuanced contextual relationships40. Moreover, PPI network databases are fragmented, and biased toward well-studied “hub” genes, limiting discovery of rare disease-relevant functional modules41,42.

Lastly, mutational signature analysis methods characterize mutagenic processes from local sequence context^43^. While invaluable for understanding tumor biology, these methods are not designed to capture interactions across multiple mutations. Furthermore, variability in sequencing platforms, where not all genes are uniformly sequenced may introduce platform-specific biases that hinder cross-cohort integration^44^. Altogether, existing approaches remain considerably limited by data sparsity, platform heterogeneity and incomplete biological priors, which hampers reliable discovery of clinically relevant mutation patterns.

Large Language Models (LLMs) have significantly transformed natural language processing by capturing semantic meaning of words through rich context-dependent vector representations or *embeddings*^45^. Just as words derive meaning from their surrounding context, the biological significance of a mutation may depend on its broader genomic context: in particular, the presence or absence of specific co-occurring mutations. Transformers offer a powerful and flexible framework for modeling such contextual relationships^46,47^. By learning context-dependent vector representations of mutations across thousands of tumors, these models can reveal hidden structures in somatic mutation data, identify synergistic mutation combinations, and improve predictions of patient outcomes^48^.

Motivated by this, we introduce OncoBERT, a language model that learns rich context-aware vector representations of somatic mutations through analysis of large genomic sequencing datasets. By mapping cancer mutation profiles onto a shared representation space, OncoBERT enables clustering and classification of tumor samples across diverse sequencing platforms, uncovering recurring mutation patterns and their association with treatment response. OncoBERT is trained in a self-supervised fashion using a large corpus of approximately 250,000 tumor samples from over 211,000 patients from the AACR-GENIE database, spanning 113 tumor types and 112 targeted sequencing panels. Leveraging OncoBERT’s embeddings, we uncover 130 distinct tumor mutational patterns and investigate their associations with response and resistance to different cancer therapies. We further investigate the biological significance of these mutational patterns by linking them to transcriptomic features of tumors, leveraging matched whole exome sequencing and bulk RNA sequencing data from The Cancer Genome Atlas (TCGA). To further stimulate research on this topic, we have made OncoBERT publicly available to the community through our repository (https://github.com/spatkar94/OncoBERT).

## Results

### Overview of OncoBERT

OncoBERT is a deep representation learning model that transforms sparse somatic mutation profiles of tumors into dense, context-aware vector embeddings. These embeddings enable clustering of tumor samples into distinct molecular subtypes, revealing recurring mutation patterns. The input data for OncoBERT are mutation profiles of tumor samples represented as binary vectors, in which each entry reflects the mutation status of a protein-coding gene: 1 = presence of at least one non-silent mutation i.e., missense, nonsense, frameshifts or indel; 0 = no mutation; * = not profiled/unknown. This tabular data is subsequently transformed into ordered gene sequences, where the relative positions of genes in each sequence reflects the mutational landscape of each sample. To determine this ordering, we model mutated genes as “heat” sources within a protein interaction network and iteratively propagate heat to neighboring genes within the network. The resulting steady-state heat distribution is used to sort genes into an ordered sequence, ranging from hot to cold. As a result, genes that are functionally related or affected by coordinated mutational processes end up positioned close to one another in the generated sequence (see Figure 1A). One key distinction of our computational framework is that we construct the protein interaction network de novo using a large pretrained protein language model (Methods) instead of relying on curated biological pathways or PPI databases. The resulting ordered sequences are treated as sentences and transformed into compact context-aware vector representations using a BERT encoder47 (Figure 1B).

**Figure 1:**
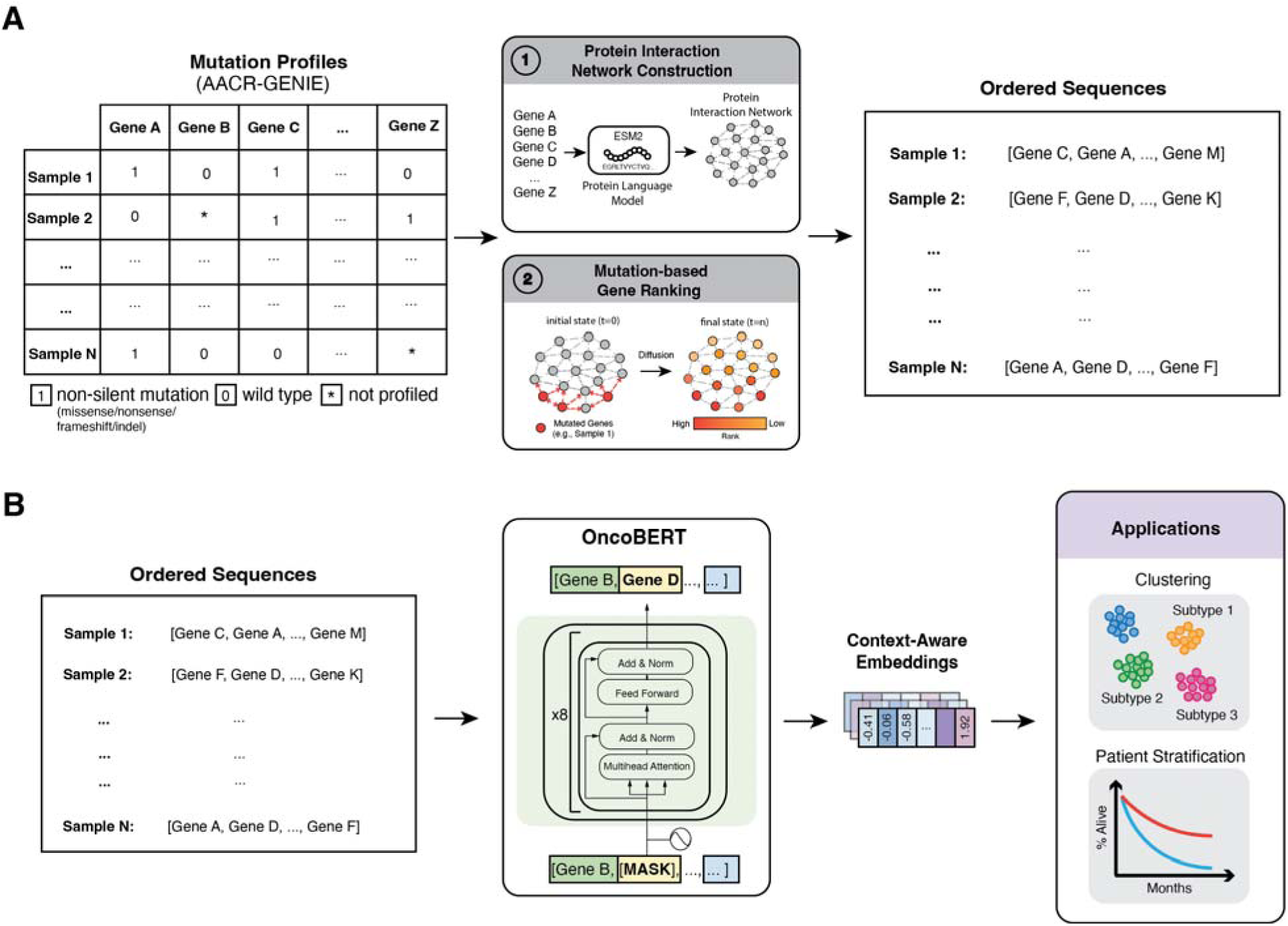
Overview of OncoBERT’s masked language modeling framework and downstream applications. **(A)** The pipeline begins with mutation profiles from the AACR GENIE database, represented as binary vectors. Here: 1 = presence of at least 1 non-silent mutation/frameshift/indel; 0 = wild type; * = gene not profiled. This data is then transformed into ordered sequences where the relative positions of genes in each sequence reflect how a specific group of mutations are organized within a protein interaction network. **(B)** These ordered sequences are treated as sentences and analyzed by OncoBERT, a transformer-based language model which transforms them into rich context aware feature embeddings. These embeddings are subsequently utilized for downstream tasks, such as unsupervised clustering and risk stratification for survival analysis.

OncoBERT is trained in a self-supervised manner using a masked language modeling (MLM) objective, where the model gradually learns to predict the identity of randomly masked genes within each sequence. This process eventually converges (Supplementary Figure 1), resulting in a model that can recognize a broad spectrum of mutation co-occurrence patterns, going beyond simple pairwise relationships. For self-supervised training, we utilized over 250,000 tumor samples from the AACR GENIE44 database; the largest publicly available real-world cancer genomics registry, spanning 113 cancer types, 112 sequencing panels, and 20 institutions. The final output of OncoBERT is a 256-dimensional embedding vector encoding the mutational patterns of each sample.

### OncoBERT Uncovers Distinct Tumor Mutation Subtypes

To investigate mutation patterns captured through OncoBERT’s embeddings, we apply a two-step unsupervised clustering approach involving Leiden community detection followed by hierarchical cluster merging^49^ (**See Methods**). This results in the discovery of 130 distinct tumor clusters or *mutation subtypes,* each characterized by a unique mutation pattern (**Figure 2A**). Furthermore, to classify new samples into one of the identified subtypes we developed a simple multi-layer perceptron (MLP) classifier that takes as input OncoBERT’s embeddings and outputs the subtype classification (**See Methods**).

**Figure 2:**
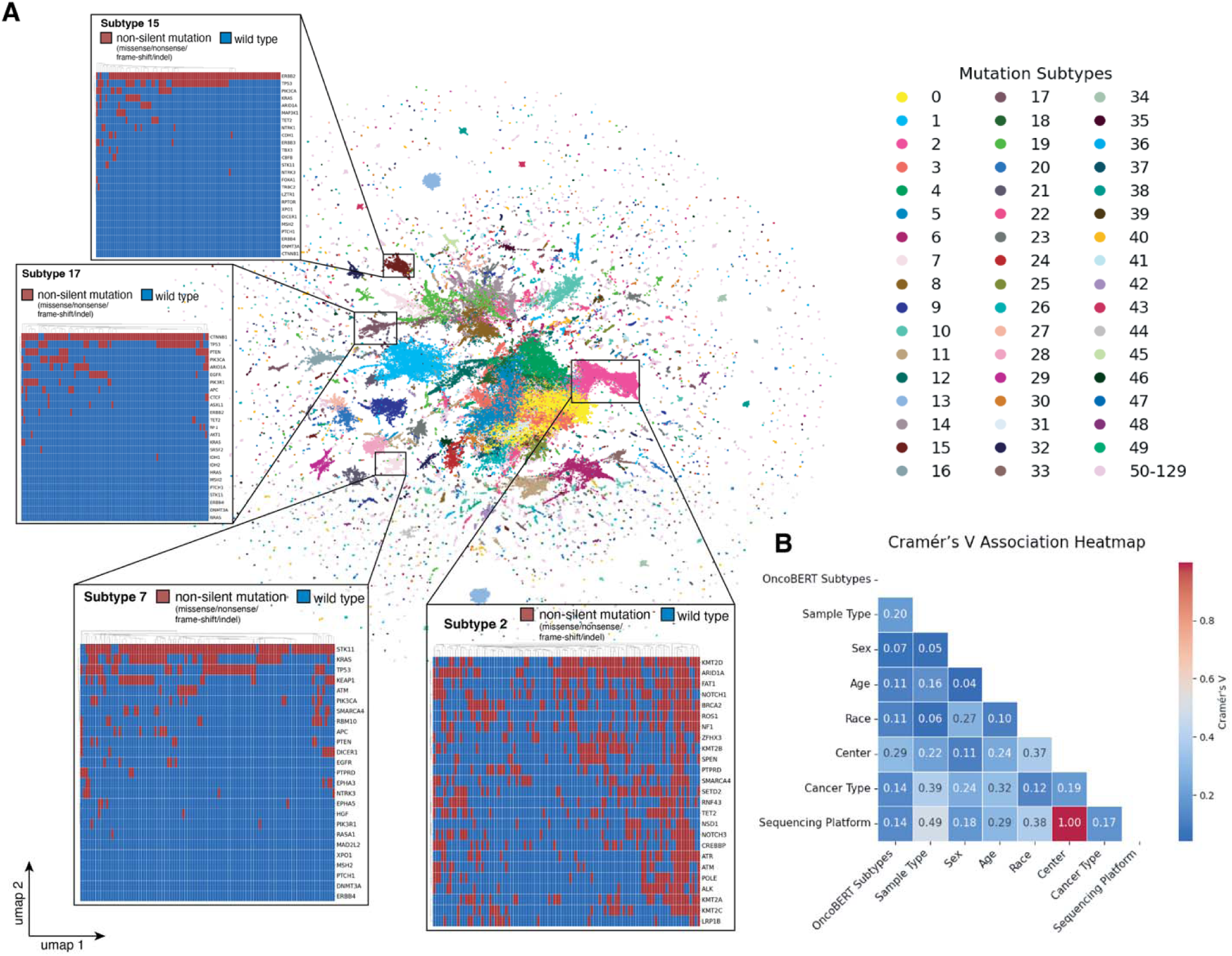
Landscape of cancer mutation patterns identified by OncoBERT. **(A)** UMAP visualization of 130 distinct genomic subtypes derived from clustering OncoBERT embeddings. Insets provide detailed mutation heatmaps for representative clusters (Subtypes 2, 7, 15, and 17), illustrating unique mutation patterns corresponding to each subtype where red indicates mutated and blue indicates wild-type/not-mutated genes. The genes depicted represent the top 25 genes contributing to a specific subtype ranked according to random forest feature importance scores **(B)** Cramér’s V association heatmap showing the correlation between OncoBERT subtypes and clinical/demographic variables, including sequencing platform, sample type, sex, age, race, center and cancer type.

To identify key co-mutated gene sets contributing to each subtype, we also trained a random forest classifier for each subtype. Specifically, each classifier was optimized to distinguish a given subtype from all others using binary mutation profiles as input features. We then leveraged the random forest classifier’s feature importance scores to rank genes most strongly associated with each subtype. This analysis uncovered distinct mutated gene sets driving each subtype. For instance, Subtype 2, highlighted in Figure 2, represents a hypermutated-subtype characterized by frequent co-occurring mutations in specific chromatin remodeling genes (ARID1A, KMT2D) as well as DNA Damage response genes (BRCA2, ATM, ATR, POLE); subtype 7 is characterized by frequent co-occurring mutations in KRAS, TP53, STK11 and KEAP1; subtype 15 is characterized by frequent co-occurring mutations in ERBB2 and TP53, and subtype 17 is characterized by frequent co-occurring mutations in CTNNB1 and TP53, along with tertiary mutations in EGFR, PI3KCA or PTEN. A detailed picture of the top-ranking genes contributing to each subtype are presented in **Supplementary Figure 2**.

To assess the degree to which OncoBERT-derived subtypes are correlated with known clinical and technical variables (e.g., sequencing platform, institution, cancer type, race, etc), we computed the Cramer’s V association between each variable and OncoBERT subtypes. Overall, we observed weak associations (range: 0.14–0.29; **Figure 2B, Supplementary Figure 3**), indicating that OncoBERT captures meaningful patterns of genetic variation, and is not confounded by known clinical and technical differences.

### Mutation Patterns Influencing Patients’ Responses to Cancer Therapy

We next investigated associations between specific OncoBERT subtypes and treatment response leveraging matched targeted sequencing and clinical outcome data from MSK-CHORD^10^, the largest publicly available database of patients with paired genomic, treatment exposure and survival outcome information. Specifically, for each drug class, each subtype was ranked based on its ability to stratify overall survival outcomes of patients, as estimated by the hazards ratio and p-value of a cancer type-stratified cox-proportional hazards model, which accounts for cancer type-specific survival differences (**Methods**). To correct for multiple hypotheses testing, all p-values were adjusted using the Benjamini-Hochberg (BH) procedure and subtypes influencing outcomes with an FDR adjusted p-value < 0.1 were classified as statistically significant.

This analysis identified subtype 2 as the strongest predictor of favorable responses to immune checkpoint inhibitors (ICI), platinum-based chemotherapeutic agents, antimetabolites, and topoisomerase inhibitors (**Figure 3A**), whereas subtype 7 consistently predicted worse outcomes across multiple therapeutic classes. Furthermore, patients harboring EGFR driver mutations and classified as subtype 17 exhibited significantly improved responses to EGFR tyrosine kinase inhibitors (TKIs), while those classified as subtype 15 demonstrated markedly poor responses. Significant associations were also observed between patient outcomes and subtypes 0,1, 8, 9, 10, 21, 49, 53, and 104. Most notably, subtype 104, which is exclusively characterized by SPOP or EAF1X mutations, exhibited superior responses to androgen deprivation therapy (ADT) in advanced prostate cancer (**Figure 3B, 3C**). In contrast, subtype 0, defined by co-occurring mutations in TP53/PTEN or CDH1/PIK3CA and the absence of KRAS mutations, was associated with significantly poor responses to ADT in prostate cancer as well as estrogen receptor–targeted hormonal therapies in breast cancer (**Figure 3D**). Subtype 21, which is enriched for co-occurring TP53/KRAS mutations as well as alterations in SMAD4 and APC, was associated with the worst outcomes in the context of ADT in prostate cancer as well as treatments involving alkylating agents (**Figure 3E**).

**Figure 3:**
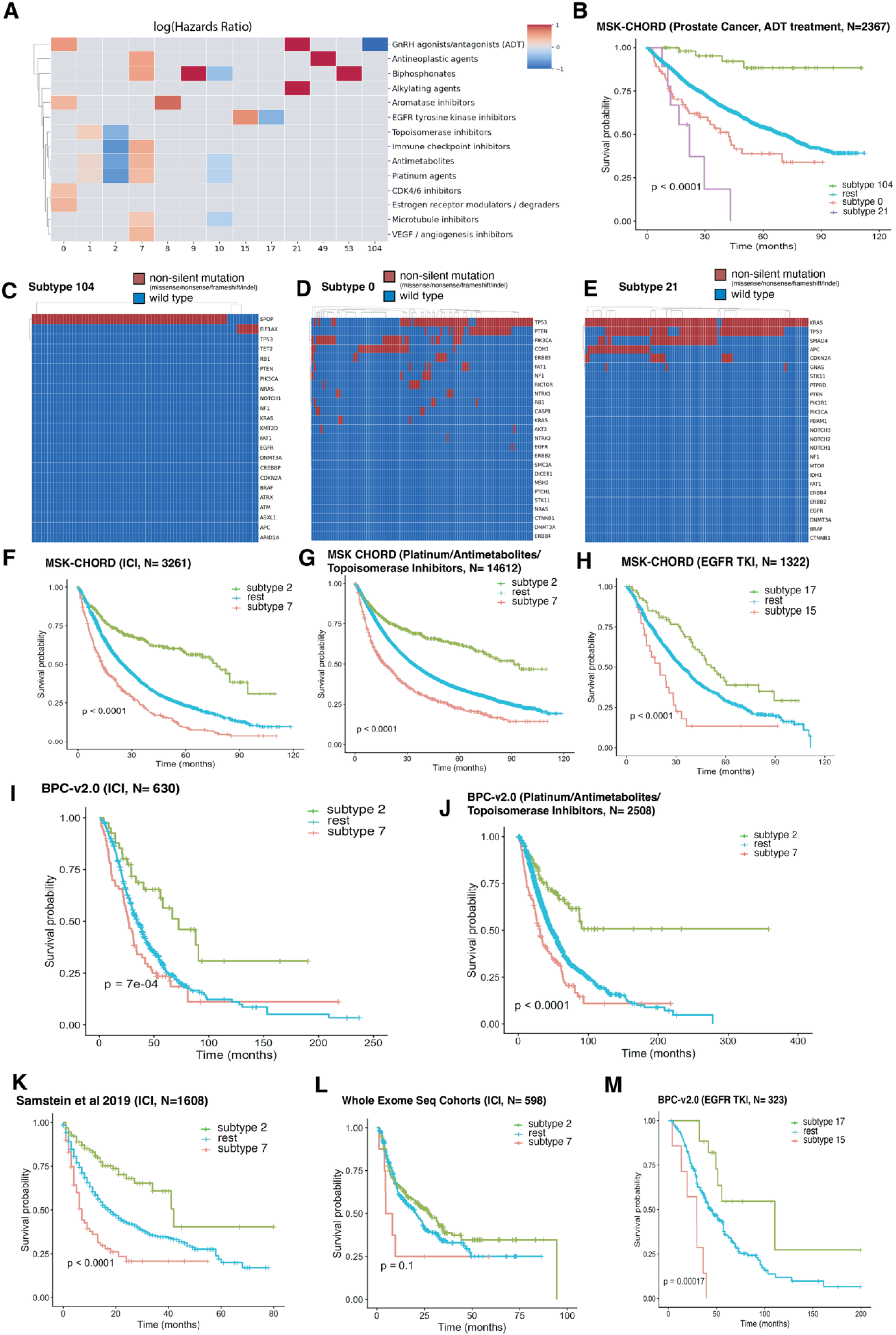
OncoBERT mutation subtypes predict differential treatment response and patient survival. **(A)** Heatmap showing the log(Hazards Ratio) for specific OncoBERT subtypes across various therapeutic classes, highlighting subtypes with significantly better (blue) or worse (red) prognosis. **(B–E)** Detailed analysis of advanced prostate cancer patients (MSK-CHORD, N=2,367) treated with Androgen Deprivation Therapy (ADT). Kaplan-Meier curves (B) show distinct survival outcomes for Subtypes 104, 0, and 21, with their corresponding mutational profiles displayed in heatmaps (C–E). **(F–H)** Kaplan Meier curves of patients from MSK-CHORD who received Immune Checkpoint Inhibitor Therapy (F), platinum-based chemo/antimetabolite/topoisomerase inhibitor therapy (G), EGFR TKI therapy (H) **(I-M)** Validation of risk stratification in patients treated with Immune Checkpoint Inhibitors (ICI), platinum-based chemo/antimetabolite/topoisomerase inhibitor therapy and EGFR TKI across independent cohorts (Samstein 2019, BPC-v2.0, and Whole Exome Seq Cohorts).

We further investigated generalizability of some of these associations in additional cohorts, wherever matched treatment exposure and overall survival information were available (**Methods**). In pan-cancer analyses, we observed consistent associations between OncoBERT subtypes 2 and 7 and patient prognosis following exposure to platinum-based chemotherapies, anti-metabolites, topoisomerase inhibitors or ICIs (**Figure 3F-G, I-L**). However, the magnitude and direction of these associations varied across different cancer types, with strongest effects observed in lung, colon and skin cancers (**Supplementary Figure 4**). Beyond ICI and chemotherapy, we were also able to replicate associations between OncoBERT-assigned mutational subtypes 15 and 17 and patient prognosis in the context of EGFR TKI therapy (**Figure 3H,M**).

### Integrating OncoBERT with Established Genomic Biomarkers Enhances Stratification of Patient Responses to Immunotherapy

We next investigated whether incorporating OncoBERT-based tumor subtypes with established genomic biomarkers, such as the overall tumor mutation burden (TMB), microsatellite instability scores (MSI), as well as specific gene/pathway-level TMB scores (NeST)^50,51^ and mutational signature activity, further enhances prediction of patient responses to immunotherapy. Hence, we fit multivariate Cox proportional hazards models to predict overall survival of patients receiving ICI therapy in MSK-CHORD, incorporating OncoBERT-derived mutation subtype labels alongside TMB, MSI, NeST scores, and mutational signature exposures (**Figure 4A**). Across different co-variates, OncoBERT-derived tumor subtypes 2 and 7 consistently emerged as significant predictors of patient survival, independent of other genomic biomarkers (**Figures 4B-D**). Importantly, across distinct clinically defined TMB and MSI subgroups, subtypes 2 and 7 effectively stratified survival outcomes (**Figure 4E-G**), highlighting their utility for more refined risk assessment and clinical decision-making for immunotherapy.

**Figure 4:**
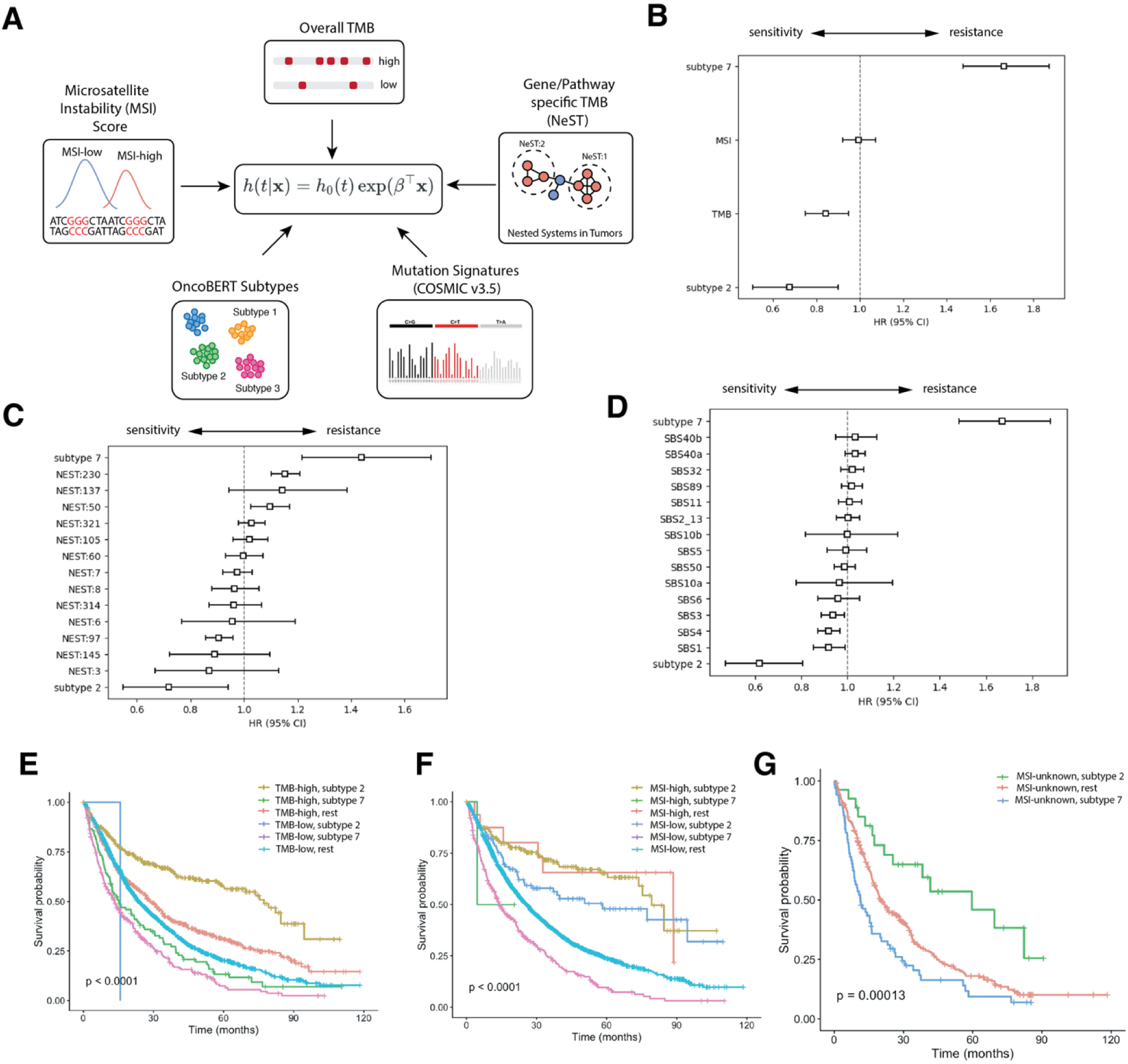
Integration of OncoBERT subtypes with established biomarkers enhances immunotherapy response stratification. **(A)** Schematic of Cox proportional hazards models incorporating Microsatellite Instability (MSI) scores and overall Tumor Mutational Burden **(B)**, gene/pathway-specific TMB scores **(C)**, and Mutation signatures **(D)** along with OncoBERT subtypes 2 and 7. **(B-D)** Forest plots displaying Hazard Ratios (HR) and 95% Confidence Intervals (CI), identifying OncoBERT subtype 7 and subtype 2 as the strongest independent predictors of poor and favorable prognosis, respectively. **(E-G)** Kaplan-Meier survival curves demonstrating the stratification power of OncoBERT subtypes across different clinically defined patient subgroups. Subtype 2 consistently identifies a superior survival group regardless of whether patients have (E) high/low TMB, (F) MSI-high/low status, or (G) unknown MSI status, while subtype 7 consistently indicates the highest risk.

Subtype 2, which was predictive of response, showed a significantly higher mutational burden across several known COSMIC signatures compared to other subtypes (p < 0.0001; **See Supplementary Figure 5**). Most notably we observed a strong enrichment of signatures linked with defective DNA repair: particularly SBS10a/b (linked to POLE exonuclease domain defects), SBS6 (associated with mismatch repair deficiency) and SBS3 (Homologous recombination deficiency), and aging (SBS1 and 3). Whereas APOBEC signatures (SBS2/13) were notably depleted. Besides endogenous signatures, Subtype 2 also showed strong enrichment for environmental signatures, such as SBS4 (tobacco smoking). Nevertheless, in a multivariate cox analysis, we see that subtype 2 remained strongly predictive of outcomes even after accounting for these associations (**Figure 4D**). These results suggest that OncoBERT’s context-aware molecular representations capture clinically relevant interactions across different mutational processes that are missed by existing biomarkers.

### Functional and Immunological Characteristics of OncoBERT-derived Tumor Subtypes

Finally, we investigated the biological significance of OncoBERT-derived tumor mutation subtypes by linking them to specific downstream transcriptional programs. Specifically, we assessed the transcriptional activity of canonical cancer hallmark processes^52,53^ within each OncoBERT subtype using matched whole-exome sequencing and bulk transcriptomic profiles from The Cancer Genome Atlas (TCGA). For each subtype, we measured the log2 fold change in enrichment scores relative to all other subtypes and estimated p-values for differential activity (**See Methods**).

Despite substantial transcriptional heterogeneity across tumors, we observed significant enrichment patterns (FDR < 0.1) with specific mutation subtypes converging on unique functional themes (**Figure 5A**). For instance, Subtype 2, was enriched for MTOR signaling, unfolded protein response, DNA repair, cell cycle and interferon response pathways, consistent with a highly proliferative and immunologically active tumor state. Subtype 7, which was frequently characterized by co-occurring KRAS, STK11 and KEAP1 mutations, was enriched for oxidative phosphorylation. This aligns with recent studies linking these concurrent mutations to metabolic rewiring and an aggressive, treatment-resistant phenotype^10,54,55^. Subtype 17, which was enriched for CTNNB1 co-mutations, exhibited strong Wnt signaling activity (log-2 fold change: 1, p-value < 0.001) and down regulation of immune response pathways^56^, whereas subtype 1, characterized by frequent co-occurring mutations in TP53, KRAS, PIK3CA and APC, was enriched for processes such as P53 signaling, TGF-β signaling, Wnt and Notch Signaling.

**Figure 5:**
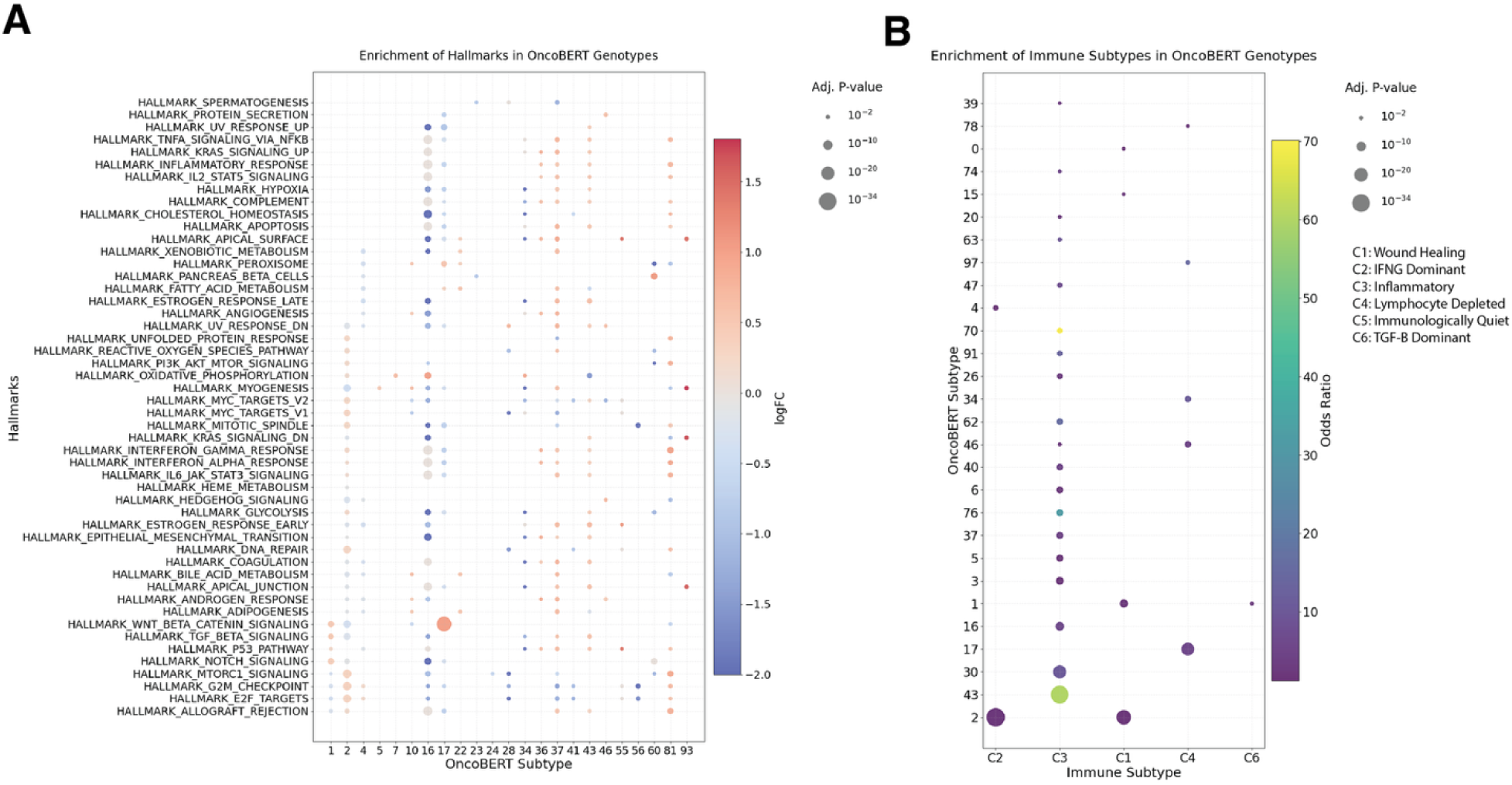
Functional characterization of OncoBERT-derived tumor subtypes. **(A)** Dot plot showing the relative enrichment of specific cancer hallmark gene sets across distinct OncoBERT-derived tumor subtypes (Only significant associations shown, i.e., FDR adjusted p-value < 0.1). Red indicates high enrichment for specific processes, while blue indicates down-regulation. **(B)** Dot plot showing the enrichment of established consensus immune microenvironment subtypes (C1–C6) with specific OncoBERT subtypes (Only significant associations shown, i.e., FDR adjusted p-value < 0.1). The size of each dot represents the FDR adjusted p-value, and the color indicates the odds ratio, with larger, brighter dots highlighting stronger associations, such as the enrichment of Subtype 43 with the C3 (Inflammatory) state and Subtype 2 with the C2 (IFNG Dominant) state.

Beyond intrinsic tumor programs, we also investigated enrichment patterns between specific OncoBERT subtypes and established pan-cancer tumor immune microenvironment (TME) subtypes^57^: C1 (Wound Healing), C2 (Interferon Gamma Dominant), C3 (Inflammatory), C4 (Lymphocyte Depleted), C5 (Immunologically Quiet), and C6 (TGF-β Dominant), (**Figure 5B**, **Methods**). Overall, we observed several significant associations (hypergeometric test FDR adjusted p-value < 0.1) indicating that specific mutational patterns exert a measurable influence on the TME. For instance, Subtype 2 showed strongest enrichment for the Interferon Gamma Dominant (C2) TME subtype (hypergeometric test, p < 0.001), consistent with tumor-intrinsic hallmark processes activated within this subtype. Subtype 43 displayed a striking enrichment for the C3 inflammatory immune subtype (hypergeometric test, FDR-adjusted p-value = 2 × 10⁻³², odds ratio = 60.12). Meanwhile, Subtype 17 was associated with a lymphocyte-depleted TME subtype (C4). Collectively, these results demonstrate that specific mutation patterns identified by OncoBERT define biologically distinct tumor states, influencing both intrinsic oncogenic programs and the tumor microenvironment.

## Discussion

In this work, we introduced OncoBERT, a language model that learns to decode tumor genetic heterogeneity through analysis of vast unlabeled genomic sequencing datasets. A key contribution of OncoBERT lies in its ability to learn robust contextual representations of somatic mutations across various sequencing platforms and centers. This is particularly valuable in the era of precision oncology, where different institutions often vary in their sequencing panel design, depth and sample collection. Moreover, from a methodological perspective, OncoBERT represents a conceptual shift in cancer genomics, moving from isolated gene and pathway-level analyses^35,50,58^, toward fully data-driven, contextual modeling of cancer mutational landscapes.

By analyzing a large cohort of over 250,000 tumor samples from AACR-GENIE, we uncovered multiple clinically relevant mutational contexts associated with response and resistance to chemotherapy, targeted therapy, and immunotherapy. Many of these associations are supported by independent clinical and experimental studies. For instance, loss-of-function mutations in KEAP1 and STK11 have been associated with an aggressive tumor phenotype and resistance to multiple therapies, including immunotherapy^10,59,60^. Recent experimental work has further shown that these tumors exhibit an altered metabolic phenotype, making them vulnerable to ATR inhibition^54^ and Ferroptosis^55^. Similarly, in locally advanced or metastatic prostate cancer, SPOP mutations have been linked to heightened androgen receptor (AR) signaling dependence and exceptional responses to androgen deprivation therapy^61,62^. In contrast, loss-of-function mutations involving tumor suppressors such as PTEN have consistently been associated with a more aggressive, treatment-resistant phenotype^63^.

A particularly interesting observation in this study is the association between specific EGFR mutation patterns and responses to EGFR TKIs. We find that EGFR-mutant tumors harboring co-occurring CTNNB1 mutations (subtype 17) exhibit improved responses to EGFR inhibitors relative to the baseline population, whereas tumors with concurrent TP53 and ERBB2 mutations (subtype 15) showed inferior responses. Although the association between EGFR-CTNNB1 co-mutations and EGFR-TKI response appears to contradict preclinical studies, which suggest a role in treatment resistance^64,65^, recent independent clinical data suggests these co-mutations are in fact beneficial for patients treated with Osimertinib therapy^66^, highlighting the complex nature of crosstalk between EGFR and Wnt signaling pathways.

In the context of immunotherapy, we find that integrating OncoBERT’s predictions with existing biomarkers significantly improved stratification of patient responses. Upon further integration with transcriptomics data, we uncovered significant associations between specific OncoBERT subtypes, cancer hallmark processes, and characteristics of the tumor microenvironment. Most notably, subtype 2, which was found to be strongly associated with immunotherapy response, preferentially activated interferon alpha and gamma signaling, which play central roles in regulating PDL1 expression and ICI response^67,68^. Among its various mutations, subtype 2 had the highest frequency of co-occurring mutations in ARID1A and KMT2D (∼32%). ARID1A is a frequently mutated tumor suppressor and crucial subunit of the SWI/SNF chromatin remodeling complex^69^, while KMT2D, is a histone methyltransferase that regulates transcription, particularly at enhancer regions^70^. While mutations in both genes have been independently studied and linked with ICI efficacy^71–74^, their combined impact within the broader mutational context is poorly understood. Our findings show that OncoBERT’s contextual representations capture clinically meaningful genetic dependencies that extend beyond these individual mutations or aggregate mutation counts, further enhancing prediction of patient outcomes. This highlights the clinical significance of considering the broader mutational context when making therapeutic decisions.

While promising, there are certain limitations of this study that can be improved upon in future work. First, although OncoBERT captures rich contextual features, it currently operates on gene-level mutation calls and does not incorporate more fine-grained mutation information such as gain or loss of function effects (e.g., AlphaMissense^75^) and variant allele frequencies^76^. Second, extending the framework to incorporate additional types of alterations, such as copy number alterations or gene fusions could further enrich learned representations^77^. Third, tumor-type–specific fine-tuning may be required to reliably capture genetic heterogeneity within individual cancer types, especially those that are typically rare or underrepresented. Fourth, the interpretability of transformer attention layers remains an open challenge. In this work, we leveraged random forest analysis to highlight recurrently co-mutated genes captured by OncoBERT’s embeddings. This analysis was inspired from previous work that leveraged random forests to characterize key methylation patterns defining different brain tumor subtypes^78^. In the future, more sophisticated methods (e.g Sparse Autoencoders) could be applied to help further decode the internal logic underlying OncoBERT’s learned latent representations^79^.

In summary, OncoBERT presents a powerful framework for modeling complex mutational patterns within cancer genomes. As genomic datasets continue to expand, OncoBERT can leverage increasing data scale to learn more robust and generalizable representations of tumor biology. By linking these learned representations to therapeutic outcomes, it has the potential to identify novel biomarkers that improve treatment selection and further strengthen precision oncology workflows^80–82^. Going beyond analysis of mutations, OncoBERT’s contextual representations can be integrated with additional patient data including medical imaging, electronic health records, spatial proteomics and digital pathology to build more comprehensive predictive models of treatment response ^83^. Such multi-modal predictive models will be key to building accurate digital twins that will ultimately guide individualized cancer care^84,85^.

## Acknowledgements

We would like to express our sincere gratitude to Dr. Alejandro Schaffer, Dr. Travis Stracker and Dr. Deborah Citrin for their time and expertise in reviewing the initial draft of this manuscript. Their insightful feedback and valuable suggestions significantly improved the quality and clarity of this work. This work was supported by the Intramural Program of the National Cancer Institute.

## Disclaimer

The content of this publication does not necessarily reflect the views or policies of the Department of Health and Human Services, nor does mention of trade names, commercial products, or organizations imply endorsement by the U.S. Government. The funders had no role in study design, data collection and analysis, decision to publish, or preparation of the manuscript.

## Methods

### 1. Datasets Curated and Analyzed in this Study

Our analyses drew on multiple real-world genomic sequencing datasets, including whole exome sequencing and targeted sequencing along with matched treatment exposure and overall survival outcome information. Together, these datasets provide substantial breadth and depth across tumor types and institutions, suitable for AI model development and validation purposes. Across all datasets considered, tumor samples with 0 non-silent mutations called (i.e, missense, nonsense, frameshift or indel mutations) were excluded from further analysis as those samples contain no relevant mutation information for further analysis. All mutation datasets are publicly available for download at (https://www.cbioportal.org/)

I. **AACR Project GENIE v18.0-public**^44^. For OncoBERT’s self-supervised training, we used the v18.0-public release of AACR Project GENIE, a large international pan-cancer registry that harmonizes somatic mutation data and demographics of cancer patients from approximately 20 participating centers. This release includes ∼250,000 tumor samples from more than 211,000 patients. GENIE v18.0 spans over 112 cancer types and subtypes as defined by OncoTree^86^. For each sample, genomic, demographic, and basic tumor-level clinical annotations (such as sequencing panel, center, sample type, patient age, cancer type, histologic subtype, race) are provided in a standardized format and made available via Cbioportal and Synapse. Sequencing of GENIE samples was performed on a heterogeneous set of platforms, depending on the contributing institution. For example, Dana-Farber samples were sequenced with three versions of its OncoPanel consisting of 275, 300, and 447 genes. Overall, 112 distinct sequencing panels were identified in this cohort highlighting the diversity of targeted next generation sequencing platforms currently utilized in real world clinical settings.
II. **GENIE Biopharma Collective:** GENIE-BPC provides detailed clinical follow-up information for selected tumor samples from the GENIE database by pairing GENIE mutation data with extensively curated real-world clinical outcomes, including progression free survival and overall survival. The publicly released GENIE-BPC cohorts included in this study are the NSCLC v2.0 dataset ^87^(n = 1,932 samples from 1846 patients across MSKCC, Dana-Farber Cancer Institute, Vanderbilt-Ingram Cancer Center, and University Health Network) and the CRC v2.0 dataset^88^ (n = 1,532 samples from 1485 patients across MSKCC, Dana-Farber, and Vanderbilt). These datasets include systemic therapy information, overall survival, pathology, and oncologist-documented progression, enabling association of specific mutations to overall survival of patients.
III. **MSK-CHORD**^10^: MSK-CHORD is a large curated real-world dataset generated at Memorial Sloan Kettering Cancer Center, linking rich clinical annotations derived from automated processing of real world clinical reports with targeted sequencing data from MSK-IMPACT, a hybrid-capture next-generation sequencing panel that, over time, has included sequencing of different gene sets (e.g., versions with 341, 410, and 468 genes).The dataset includes sequenced tumors from 24,950 patients, representing a broad distribution of common and rare cancers. Clinical annotations incorporate both structured fields and natural-language–extracted variables, enabling detailed analyses of treatment histories, response assessments, and disease trajectories. Major tumor cohorts include non–small-cell lung cancer (∼7,809 cases), breast cancer (∼5,368 cases), colorectal cancer (∼5,543 cases), prostate cancer (∼3,211 cases), and pancreatic cancer (∼3,109 cases).
IV. **Samstein et al, 2019, Immune checkpoint inhibitor cohort**^89^: This dataset consists of tumor samples from 1662 patients spanning diverse cancer types treated with anti–PD-1/PD-L1, anti–CTLA-4, or combination immunotherapy. For each patient, matched MSK-IMPACT targeted sequencing and overall survival information was available and utilized for survival analysis, linking specific mutation patterns to outcomes. All data is publicly available for download on Cbioportal.
V. **The Cancer Genome Atlas (TCGA)**^90^: For functional and immunogenomic analyses, we used matched whole-exome sequencing and bulk RNA-sequencing data from The Cancer Genome Atlas (TCGA), comprising approximately 10,000 tumors across 33 cancer types. All data is publicly available for download on the Genomic Data Commons (GDC) web portal (https://portal.gdc.cancer.gov/). Pan cancer Tumor microenvironment subtype annotations were derived from supplementary material of a prior pan cancer immunogenomic study on the tumor microenvironment (Thorsson et al, 2018: Immune Landscape of Cancer).
VI. **Immune checkpoint inhibitor treated whole exome sequencing cohorts:** We additionally analyzed 598 tumor samples from eight different whole-exome sequencing (WES) cohorts consisting of patients with advanced or metastatic cancers and treated with immune checkpoint inhibitors (anti-PD1/PD-L1, anti-CTLA4 or combination), each with matched overall survival outcomes. These included three cohorts from metastatic melanoma (Snyder et al.; Van Allen et al.; Hugo et al.)^91–93^, one from clear cell renal cell carcinoma (Miao et al.)^94^, two from non–small-cell lung cancer (Rizvi et al.; Hellmann et al.)^95,96^, one from glioblastoma (Zhao et al.)^97^, and one microsatellite-stable mixed solid tumors cohort^98^. All whole exome sequencing data and clinical annotations from these cohorts have been harmonized and made accessible via Cbioportal (Immunogenomic studies section).

### 2. OncoBERT Model

OncoBERT is an encoder-only transformer^47^ with a bi-directional self-attention architecture, developed to learn dense, context-dependent representations of sparse somatic mutation profiles. The model takes as input the non-synonymous mutation profile of a tumor sample, derived from either targeted next-generation sequencing (NGS) panels or whole-exome sequencing (WES) data. We focus specifically on non-synonymous mutations i.e., missense, nonsense, frameshift and indels, as these alter the protein-coding sequence and are more likely to impact protein function, either through gain-of-function or loss-of-function effects^99^.

#### I.#Representation of Tumor Mutation Profiles

For modeling purposes, we represent each tumor’s mutation profile as a binary vector:

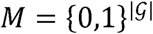

where *G*, is the set of all cancer associated protein-coding genes profiled across different sequencing platforms^100^. A value of 1 indicates presence of at least one non-silent mutation (i.e., missense, nonsense, frameshift or indel), whereas a value of 0 indicates absence. Furthermore, we represent each protein-coding gene using embeddings from a pretrained protein language model (pLM): ESM2^101^, which provides a vector representation for each gene:

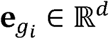

These embeddings capture diverse protein properties, including structural, evolutionary, and functional characteristics, which serve as informative priors for downstream functional modeling tasks. Using the pretrained pLM model, we construct a de novo protein interaction network *G = (V, E)*, where nodes correspond to protein-coding genes and edges represent interactions between each pair of genes based on cosine similarity of their pLM embeddings:

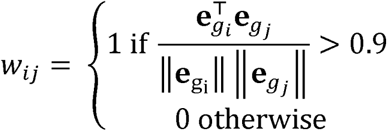

The normalized adjacency matrix **Ф** *∊ R^|G|G^*, representing the network is defined as follows:

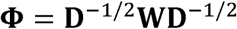

where:

- **W=[***w_ij_*] is the adjacency matrix
- **D** *∊ R^|G|G^* is the diagonal degree matrix with *D_ij_ ∑_j_w_ij_*

#### II.#Capturing the Functional Context of Cancer Mutations

We next capture the functional context of mutations by performing a random walk with restart^102^ on the protein interaction network *G*. Let **r**_0_ *∊ R^|G|^* be the initial state vector:

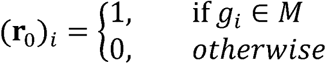

Here, M ⊂ G, represents the set of all genes harboring at least one non-synonymous mutation. Using the normalized adjacency matrix **Φ**, we iteratively compute the next state:

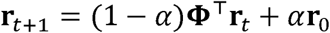

with *α* = 0.5 represents the random walk restart probability. We run this iterative process until we reach a steady state **r_*_**, which reflects the proximity of each gene to all mutated genes in the protein interaction network *G*. All genes are then ranked based on their steady state values, producing a unique ordered sequence:

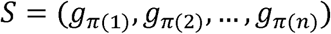

where *π,* is an ordering such that (**r**_*_)_π_ ≥ (**r**_*_)_π(2)_ ≥…. This ordering captures the functional organization of all mutated genes within the network such that mutations targeting functionally similar genes are likely to be positioned close to each other in the sequence.

#### III.#Input Tokenization and the Transformer Encoder

Each ordered sequence *S* is treated as a sentence and each gene *g_i_* in the sequence acts as a token with its associated protein language model embedding **e***_gi_* ∊ R^d^. These embeddings are projected onto a 256-dimensional token embedding space:

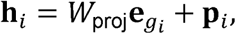

where *W*_proj_ is a linear projection matrix and **p***_i_* is a learnable positional embedding. To ensure uniform input representation across samples, each sequence is truncated to a predetermined maximum context window length *L*. These truncated sequences are then passed to a BERT-style Transformer model, which employs stacked multi-headed self-attention (MHSA) blocks with residual skip connections to learn a latent representation of each sequence:

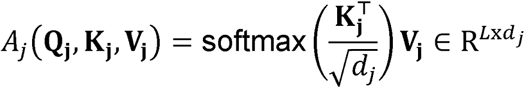

**Q_i_, K_j_, V_j_** are learnable linear projections of input token embeddings for a given MHSA block and *d_j_* = 256/(number of attention heads). The outputs of each attention head *A_j_* are concatenated and passed through another linear layer followed by layer-wise normalization to generate new token embeddings for the next block. This mechanism allows the model to capture a wide range of mutation co-occurrence patterns, including higher-order interactions and conditional dependencies, that are otherwise missed by traditional matrix factorization methods. The final latent representation generated by the transformer is the output of the class token: **z** ∊ R^256^ appended to the beginning of each sequence.

#### IV.#Self-Supervised Training via Masked Language Modeling

OncoBERT is trained using a masked language modeling (MLM) objective in which 20% of the tokens in each training batch are randomly masked. Let *M* denote the set of masked token positions. The model is then trained to predict the identity of each masked token using a binary cross-entropy loss:

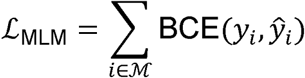

*y_i_* ∊ {0,1}^|*G*|^ represents the identity of each masked token, and *ŷ_i_* ∊ [0,1]^|*G*|^ is the predicted probability distribution. This self-supervised setup allows the model to learn a rich, context-dependent feature representation of each sample without relying on any labeled data.

#### V.#OncoBERT Architecture and Training Hyperparameters

The OncoBERT architecture consists of 8 stacked MHSA blocks, with maximum sequence length *L* = 50 with 8 attention heads per block. Self-supervised training of OncoBERT was conducted using an Adam Optimizer with a batch size of 64 and 500 epochs using mutation profiles from the GENIE v18 public dataset. Training was conducted on a single NVIDIA A100 GPU server using a learning rate of 1×10^-4^ and weight decay of 1×10^-4^ and first and second moment decay hyperparameters *β*_1_ = 0.9 and *β*_2_ = 0.999.

### 3. Genomic Subtype Discovery and Tumor Classification

Having mapped each mutation profile to a compact vector representation using OncoBERT, we first applied the Leiden clustering algorithm to cluster tumor samples into distinct subgroups^49^. Leiden clustering is an advanced graph-based clustering algorithm originally developed for high-dimensional single-cell data analysis, but broadly applicable to any dataset with dense features and complex nonlinear structure. Clustering was performed using the scanpy.tl.phenograph implementation with default parameters (k = 30, Euclidean distance metric, minimum cluster size = 10, tolerance = 0.001, maximum iterations = 2,000, and Leiden as the backend community detection method). The resulting clusters were subsequently refined using the scanpy.tl.dendogram algorithm, a bottom-up hierarchical clustering approach, in which closely related clusters were iteratively merged to construct a dendrogram. Final genomic subtype assignments were determined by conducting a simple grid search over potential dendogram height cutoffs (e.g., 0.0,0.25,0.5,0.75 and 1) and selecting the cutoff that maximized the silhouette score. This eventually resulted in the selection of a dendogram height cutoff of 0.25, resulting in the discovery of 130 distinct clusters (mutation subtypes). To assign any new tumor samples to the identified mutation subtypes, we trained a multilayer perceptron (MLP) network consisting of an input layer of 256 neurons, a hidden layer of 256 neurons, a ReLU activation with dropout probability 0.1 and a multi-class classification head. For training the MLP, we utilized the AACR-GENIE cohort, holding out 20% of the data for testing purposes. The Adam optimizer was used for updating the MLP weights with learning rate of 1×10^-3^. The trained MLP, achieved a high subtype classification accuracy on held-out test data (Precision: 0.95, Recall: 0.95, F1 score:0.95)

### 4. Association of OncoBERT Subtypes with Treatment-Specific Survival Outcomes

We evaluated associations between OncoBERT-defined genomic subtypes and treatment response using matched targeted sequencing, treatment exposure, and clinical outcome data from MSK-CHORD, the largest publicly available cohort with paired genomic and longitudinal survival information. Analyses were restricted to patients with documented exposure to a given drug class and available overall survival data.

For each drug class, we assessed the prognostic relevance of each OncoBERT subtype by fitting a cancer type–stratified Cox proportional hazards model. Let *T_i_* denote the observed overall survival time for patient *i* and let *δ_i_* indicate the event status (death vs. censoring). The hazard function for patient *i* at time *t* was modeled as:

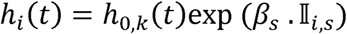

where *h*_0,*k*_ (*t*) is the baseline hazard function specific to cancer type *k*, I*_i,s_* is a binary indicator denoting membership of patient *i* in OncoBERT subtype *s*, and *β_s_* represents the log-hazard ratio associated with subtype *s*. Stratification by cancer type allows the baseline hazard to vary across tumor lineages, thereby accounting for systematic differences in survival distributions between cancer types while estimating a shared subtype-specific effect.

For each subtype–drug class pair, the estimated hazard ratio was computed as: HR*_s_* = exp (*β_s_*), and statistical significance was assessed using the Wald test derived from the partial likelihood of the Cox model. Subtypes were ranked within each drug class according to the magnitude of their hazard ratios and corresponding p-values, reflecting their ability to stratify overall survival outcomes. To account for multiple hypothesis testing across all subtype-drug class comparisons, raw p-values were adjusted using the Benjamini–Hochberg procedure. OncoBERT subtypes with a false discovery rate (FDR)–adjusted p-value below 0.1 were considered to be significantly associated with treatment-specific survival outcomes.

### 5. Calculation of NeST Scores and Mutation Signature Activity from Targeted Sequencing

NeST scores were derived from MSK-CHORD’s targeted sequencing data by projecting somatic mutations onto a hierarchical protein interaction network comprising 395 protein systems, referred to as Nested Systems in Tumors (NeST)^50^. These systems represent recurrently mutated functional units across cancer types spanning individual protein coding genes, canonical cancer pathways, and previously uncharacterized protein assemblies in which mutations converge more frequently than expected by chance. In this study, we focused on the mutation burden of 13 NeST systems previously shown to modulate response to immune checkpoint inhibitors^51^. In parallel, we estimated mutational signature activity for each tumor sample within MSK-CHORD using SATS (Signature Analyzer for Targeted Sequencing), a computational framework designed to infer robust mutational signature activity from tumors sequenced with targeted gene panels (https://github.com/binzhulab/SATS/tree/main)^103^. SATS performs de novo signature extraction using signeR^104^, maps inferred signatures to reference COSMIC single base substitution signatures^43^, and estimates sample specific signature activities and burdens. For this study, we utilized SATS to quantify the burden of known COSMIC v3.5 signatures prevalent in the cancer types represented within the MSK CHORD cohort: SBS1, SBS2, SBS13, SBS3, SBS4, SBS5, SBS6, SBS10a, SBS10b, SBS11, SBS32, SBS40a, SBS40b, SBS50, and SBS89.

### 6. Cancer Hallmark Enrichment Analysis

To characterize the functional landscape of OncoBERT-defined tumor subtypes, we computed sample-specific gene set enrichment analysis (ssGSEA) scores for 50 canonical cancer hallmark gene sets from the MSigDB Hallmark Collection^53^ across 9215 TCGA tumor samples with matched whole-exome sequencing and bulk transcriptomic data following the implementation from the following pan-cancer study^105^. Let *s_i,h_* denote the ssGSEA enrichment score of a hallmark gene-set ℎ in tumor sample *i*. For each genomic subtype *C*, we calculate the mean enrichment score for each hallmark ℎ across all samples assigned to subtype *C* as follows:

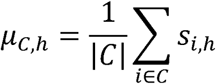

We then calculate the log fold change in mean enrichment scores between patients belonging to subtype *C* and all other subtypes *C*’ as follows:

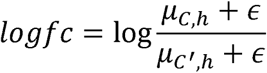

Where, *ɛ* is a small constant to prevent division by 0. By construction logfc values > 0 indicate strong positive enrichment of a given hallmark process within subtype *C* relative to other subtypes and values < 0 indicate negative enrichment. To calculate significance p-value of observed enrichments, we perform a mann-whitney U test, assessing differential activity of enrichment scores between a given subtype and all other subtypes. To account for multiple hypothesis testing across all Hallmark–OncoBERT subtype pairs, p-values were adjusted using the Benjamini–Hochberg (BH) procedure, and pairs yielding a false discovery rate (FDR) adjusted p-value < 0.1 were considered to be statistically significant.

### 7. Enrichment analysis of OncoBERT subtypes and tumor microenvironment subtypes

Given pan-cancer tumor immune microenvironment (TME) subtype annotations from TCGA (C1–C6) and OncoBERT subtype classifications, we quantified the association between each OncoBERT subtype and each TME subtype using a hypergeometric enrichment test. Specifically, for a given OncoBERT subtype–TME subtype pair, we constructed a 2×2 contingency table with the following counts:

- *a* = number of samples of TME subtype of interest within given OncoBERT subtype
- *b* = number of samples of all other TME subtypes within given OncoBERT subtype
- *c* = number of samples of TME subtype of interest in all other OncoBERT subtypes
- *d* = number of samples of all other TME subtypes in all other OncoBERT subtypes

Let *N* = *a* + *b*+ *c* +*d* denote the total number of samples, *K* = *a* + *c* be the total number of samples annotated with the TME subtype of interest, and *n* = *a* + *b* be the total number of samples in the given OncoBERT subtype. Under the null hypothesis of no association, the probability of observing exactly α overlapping samples is given by the hypergeometric distribution:

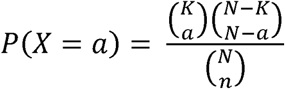

Enrichment p-values were computed as the upper-tail probability:

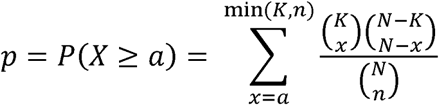

To account for multiple hypothesis testing across all OncoBERT–TME subtype pairs, p-values were adjusted using the Benjamini–Hochberg (BH) procedure, and pairs yielding a false discovery rate (FDR) adjusted p-value < 0.1 were considered to be statistically significant.

## Supplementary Figures

**Supplementary Figure 1:**
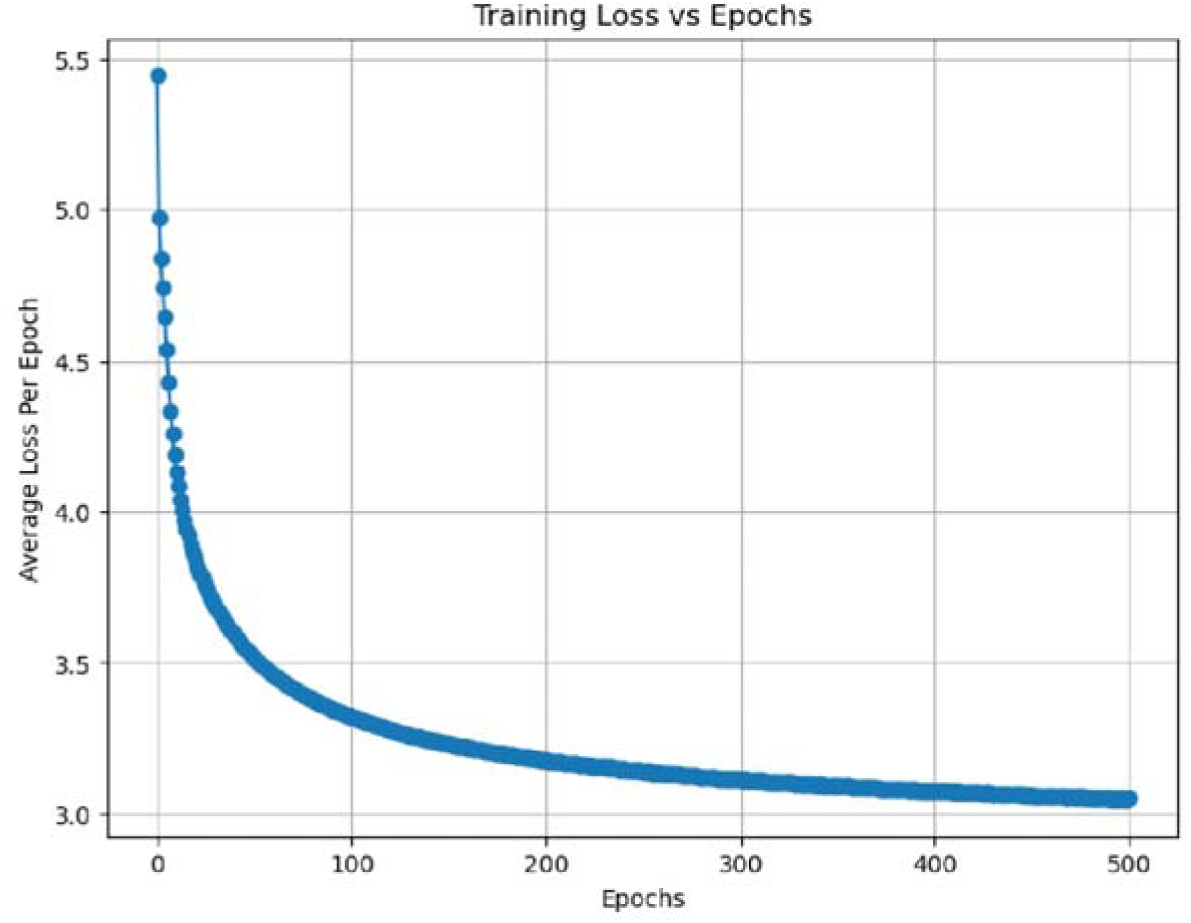
Training Loss Curve for OncoBERT. This figure shows the relationship between training duration (epochs) and error reduction during self-supervised training. The steady decline and eventual flattening of the curve suggest that the model has reached a point of diminished returns by epoch 500, confirming effective training.

**Supplementary Figure 2:**
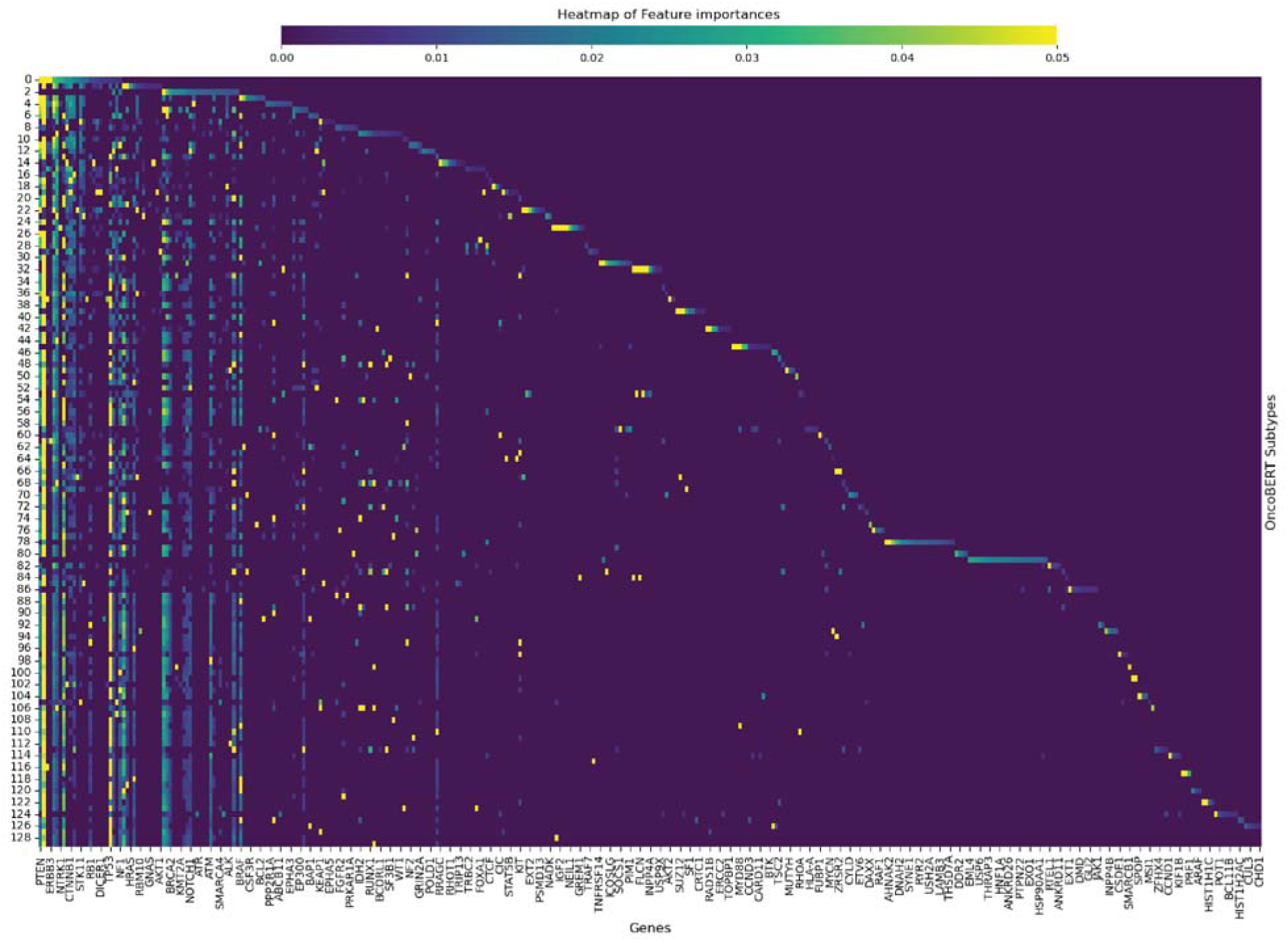
Heatmap of Genomic Feature Importance across OncoBERT Subtypes. This visualization displays the top 25 most influential genes (x-axis) for each identified OncoBERT subtype (y-axis). Feature importances were determined by random forest classifiers trained to distinguish each subtype from remaining subtypes. The color gradient indicates feature importance, ranging from 0.00 (dark purple) to 0.05 and above (yellow). The distinct diagonal pattern highlights genes that serve as primary contributors for specific subtypes, while vertical columns (e.g., PTEN, TP53) indicate frequently mutated genes with broad importance across multiple OncoBERT subtypes.

**Supplementary Figure 3:**
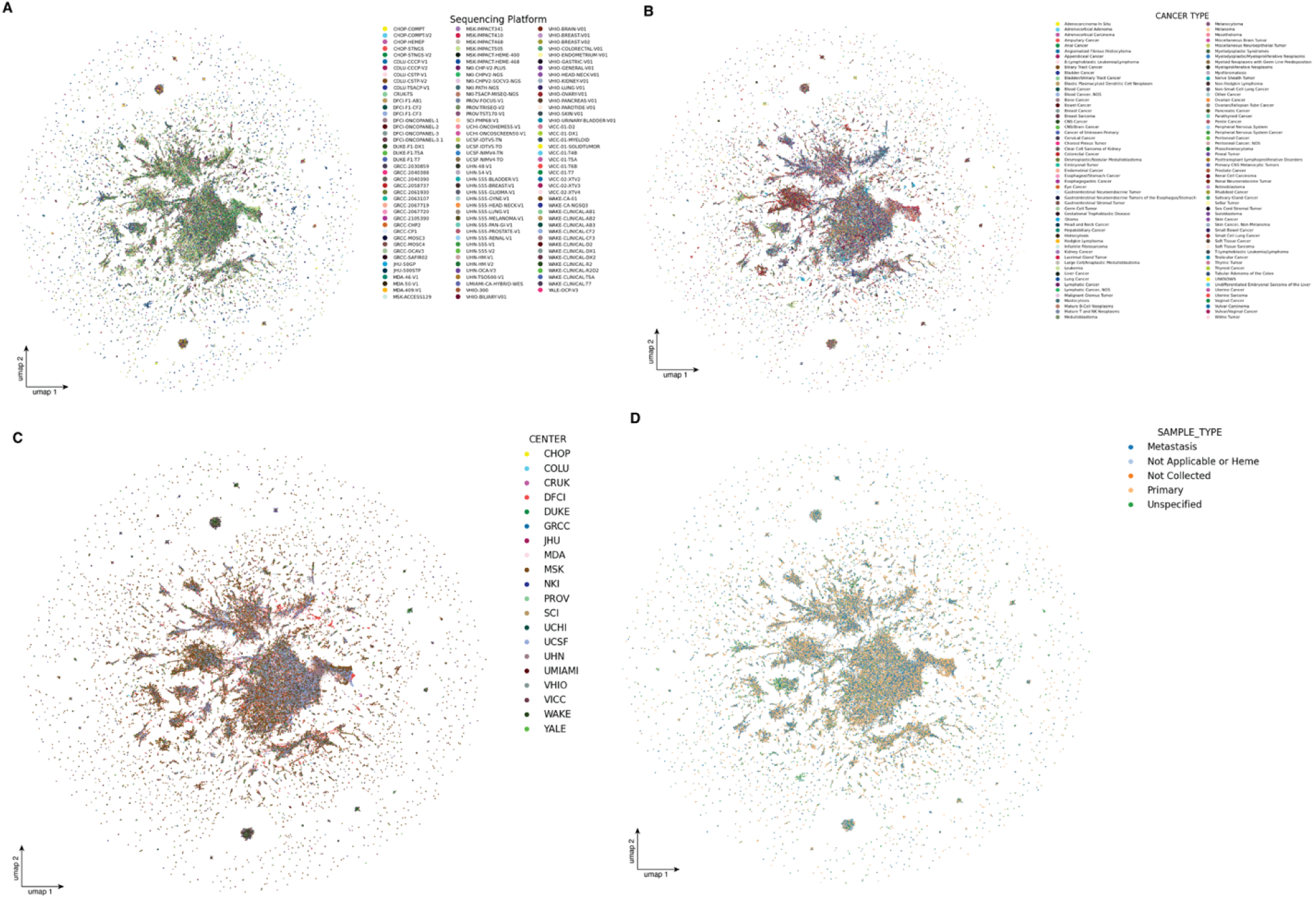
UMAP Visualization of OncoBERT Embeddings for AACR GENIE Samples. The four panels illustrate the distribution of AACR GENIE samples in OncoBERT’s embedding space colored by: **(A)** Sequencing Platform, **(B)** Cancer Type, **(C)** Sequencing Center, and **(D)** Sample Type (e.g., Primary vs. Metastasis). Samples show considerable mixing across different sequencing platforms (A) and contributing centers (B), suggesting that OncoBERT embeddings primarily capture biological signals related to malignancy rather than batch effects or technical variations.

**Supplementary Figure 4:**
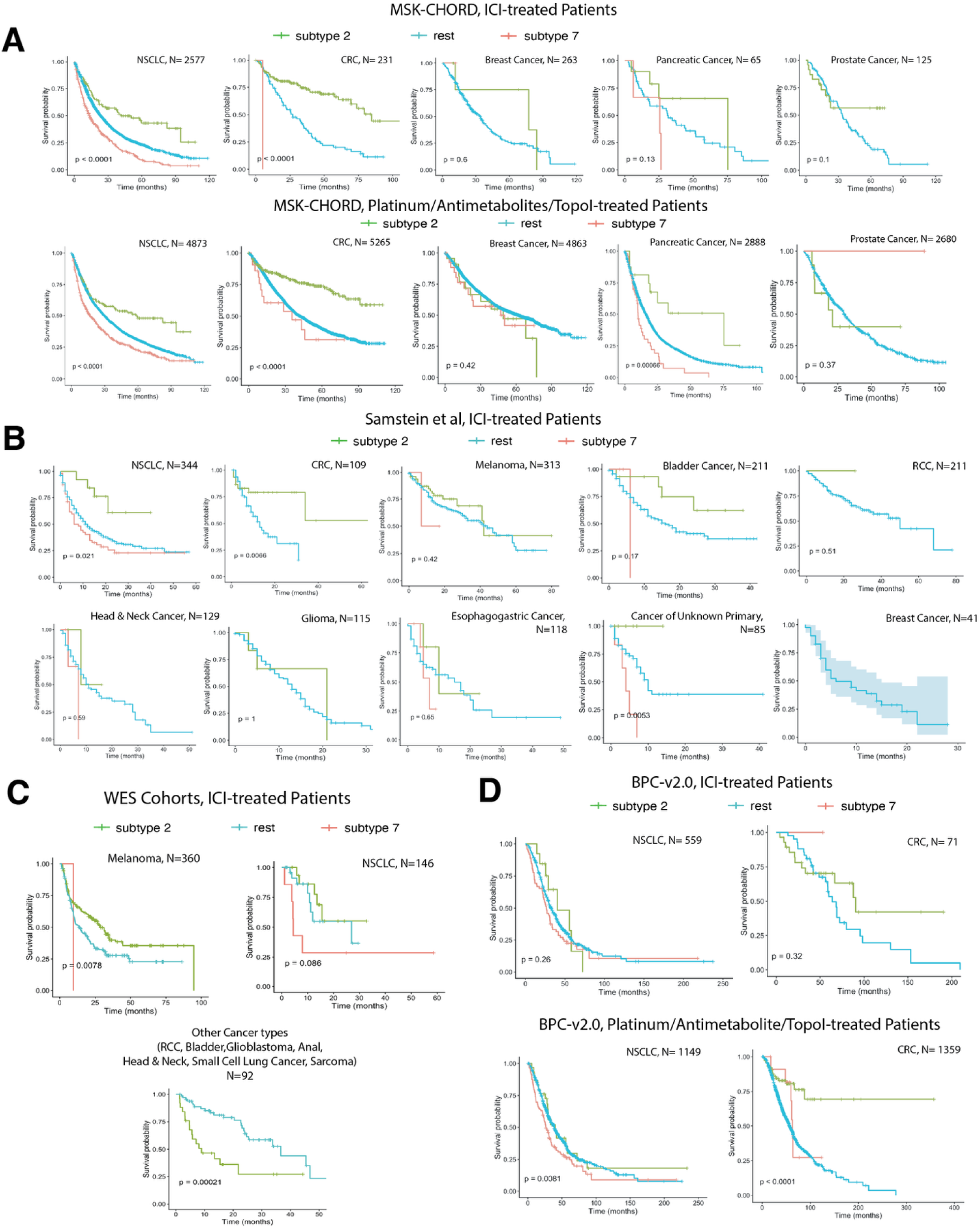
Cancer type and cohort specific differences in survival outcomes of OncoBERT-drived tumor subtypes. Kaplan-Meier analysis demonstrating the prognostic and predictive value of OncoBERT-derived classifications across multiple clinical datasets. **(A)** Analysis of the MSK-CHORD cohort, comparing Immune Checkpoint Inhibitor (ICI) therapy with cytotoxic chemotherapy (Platinum/Antimetabolites/Topoisomerase Inhibitors) across major cancer types. **(B–C)** Validation in independent ICI-treated populations, including the Samstein et al. and Whole Exome Sequencing cohorts, highlighting significant survival separation in NSCLC, Melanoma and CRC (p < 0.05). **(D)** Survival stratification within the BPC-v2.0 dataset. Across all panels, N represents the sample size per malignancy, and p-values denote statistical significance based on the log-rank test.

**Supplementary Figure 5.**
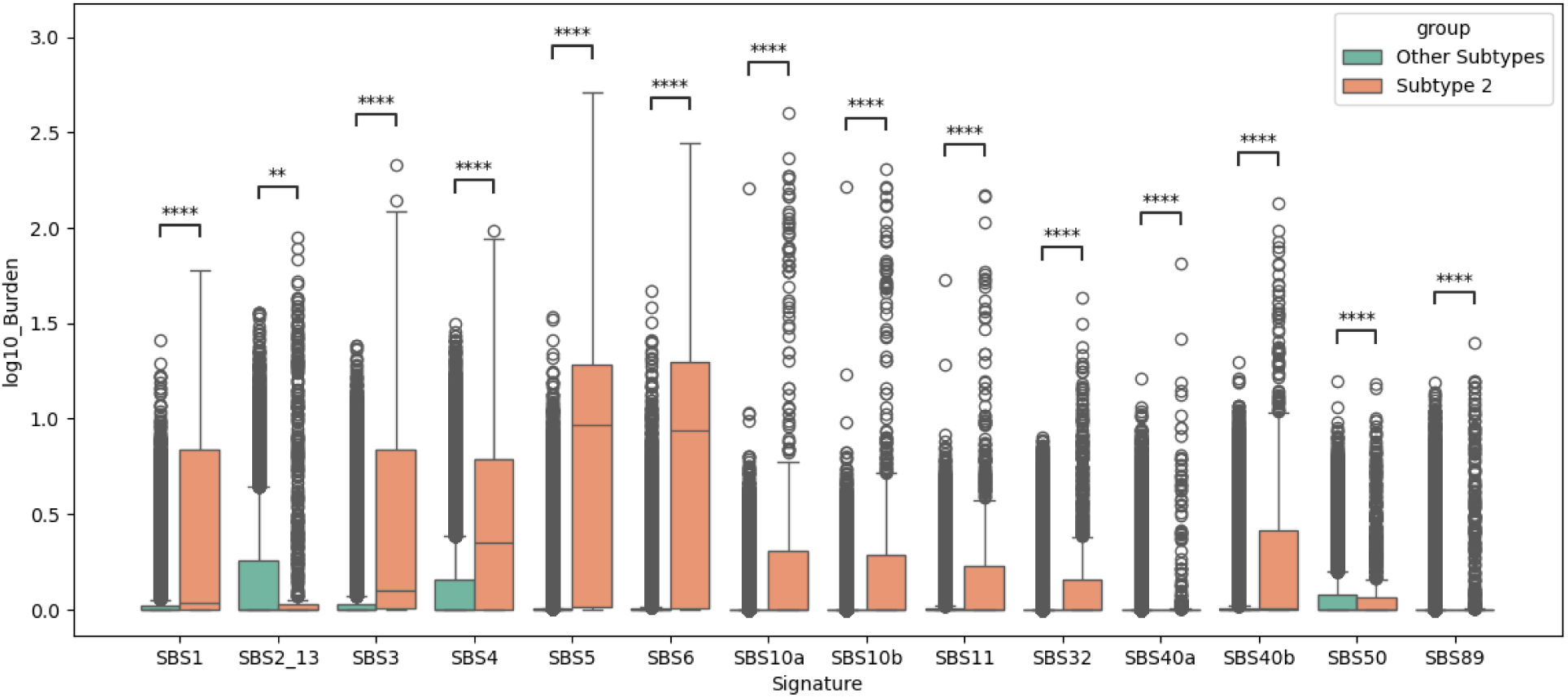
Comparison of mutational signature activity between Subtype 2 and Other Subtypes. Boxplots represent the distribution of log10 mutation burden for COSMIC Single Base Substitution (SBS) signatures across different tumor subpopulations within MSK-CHORD. Subtype 2 (orange) exhibits a significant enrichment (p-value < 0.0001) in signatures associated with DNA replication and repair deficiencies, most notably SBS10a/b (POLE exonuclease domain mutations), SBS6 (mismatch repair deficiency), and SBS3 (homologous recombination deficiency). Conversely, Subtype 2 shows a significant relative depletion in SBS2_13 activity, suggesting that APOBEC-mediated mutagenesis is a more prominent driver in Other Subtypes (green). Statistical significance was determined by Mann-Whitney U test; **p< 0.01, ****p< 0.0001.

## References

1 Black, J. R. M. & McGranahan, N. Genetic and non-genetic clonal diversity in cancer evolution. Nat Rev Cancer 21, 379–392 (2021). 10.1038/s41568-021-00336-2

2 Zahir, N., Sun, R., Gallahan, D., Gatenby, R. A. & Curtis, C. Characterizing the ecological and evolutionary dynamics of cancer. Nat Genet 52, 759–767 (2020). 10.1038/s41588-020-0668-4

3 Cheng, D. T. et al. Memorial Sloan Kettering-Integrated Mutation Profiling of Actionable Cancer Targets (MSK-IMPACT): A Hybridization Capture-Based Next-Generation Sequencing Clinical Assay for Solid Tumor Molecular Oncology. J Mol Diagn 17, 251–264 (2015). 10.1016/j.jmoldx.2014.12.006

4 Schneider, D. et al. Subtype-Specific Patterns of Tumor Purity and Mutation Load Suggest Treatment Implications: A Cross-Sectional Analysis of 7494 Soft Tissue and Bone Sarcomas (MSK Cohort). Am J Clin Oncol 48, 185–192 (2025). 10.1097/COC.0000000000001161

5 Zehir, A. et al. Mutational landscape of metastatic cancer revealed from prospective clinical sequencing of 10,000 patients. Nat Med 23, 703–713 (2017). 10.1038/nm.4333

6 Attalla, K. et al. Prevalence and Landscape of Actionable Genomic Alterations in Renal Cell Carcinoma. Clin Cancer Res 27, 5595–5606 (2021). 10.1158/1078-0432.CCR-20-4058

7 Friedman, C. F. et al. Assessing the Genomic Landscape of Cervical Cancers: Clinical Opportunities and Therapeutic Targets. Clin Cancer Res 29, 4660–4668 (2023). 10.1158/1078-0432.CCR-23-1078

8 Schrader, K. A. et al. Germline Variants in Targeted Tumor Sequencing Using Matched Normal DNA. JAMA Oncol 2, 104–111 (2016). 10.1001/jamaoncol.2015.5208

9 Nguyen, B. et al. Genomic characterization of metastatic patterns from prospective clinical sequencing of 25,000 patients. Cell 185, 563–575 e511 (2022). 10.1016/j.cell.2022.01.003

10 Jee, J. et al. Automated real-world data integration improves cancer outcome prediction. Nature 636, 728–736 (2024). 10.1038/s41586-024-08167-5

11 Carr, T. H. et al. Defining actionable mutations for oncology therapeutic development. Nat Rev Cancer 16, 319–329 (2016). 10.1038/nrc.2016.35

12 Andre, F. et al. Prioritizing targets for precision cancer medicine. Ann Oncol 25, 2295–2303 (2014). 10.1093/annonc/mdu478

13 Waarts, M. R., Stonestrom, A. J., Park, Y. C. & Levine, R. L. Targeting mutations in cancer. J Clin Invest 132 (2022). 10.1172/JCI154943

14 Schwaederle, M. et al. Detection rate of actionable mutations in diverse cancers using a biopsy-free (blood) circulating tumor cell DNA assay. Oncotarget 7, 9707–9717 (2016). 10.18632/oncotarget.7110

15 Lang, G. T. et al. Characterization of the genomic landscape and actionable mutations in Chinese breast cancers by clinical sequencing. Nat Commun 11, 5679 (2020). 10.1038/s41467-020-19342-3

16 Suehnholz, S. P. et al. Quantifying the Expanding Landscape of Clinical Actionability for Patients with Cancer. Cancer Discov 14, 49–65 (2024). 10.1158/2159-8290.CD-23-0467

17 Flaherty, K. T. et al. Molecular Landscape and Actionable Alterations in a Genomically Guided Cancer Clinical Trial: National Cancer Institute Molecular Analysis for Therapy Choice (NCI-MATCH). J Clin Oncol 38, 3883–3894 (2020). 10.1200/JCO.19.03010

18 Zill, O. A. et al. The Landscape of Actionable Genomic Alterations in Cell-Free Circulating Tumor DNA from 21,807 Advanced Cancer Patients. Clin Cancer Res 24, 3528–3538 (2018). 10.1158/1078-0432.CCR-17-3837

19 Jiang, L., Yu, H., Tang, J. & Guo, Y. CoMutDB: the landscape of somatic mutation co-occurrence in cancers. Bioinformatics 39 (2023). 10.1093/bioinformatics/btac725

20 Jiang, L. et al. Comprehensive Analysis of Co-Mutations Identifies Cooperating Mechanisms of Tumorigenesis. Cancers (Basel) 14 (2022). 10.3390/cancers14020415

21 Liu, C. et al. Individualized genetic network analysis reveals new therapeutic vulnerabilities in 6,700 cancer genomes. PLoS Comput Biol 16, e1007701 (2020). 10.1371/journal.pcbi.1007701

22 El Tekle, G., et al. Co-occurrence and mutual exclusivity: what cross-cancer mutation patterns can tell us. Trends Cancer 7, 823–836 (2021). 10.1016/j.trecan.2021.04.009

23 Dao, P. et al. BeWith: A Between-Within method to discover relationships between cancer modules via integrated analysis of mutual exclusivity, co-occurrence and functional interactions. PLoS Comput Biol 13, e1005695 (2017). 10.1371/journal.pcbi.1005695

24 Wang, X., Fu, A. Q., McNerney, M. E. & White, K. P. Widespread genetic epistasis among cancer genes. Nat Commun 5, 4828 (2014). 10.1038/ncomms5828

25 Liu, J., Zhao, D. & Fan, R. Shared and unique mutational gene co-occurrences in cancers. Biochem Biophys Res Commun 465, 777–783 (2015). 10.1016/j.bbrc.2015.08.086

26 Sinkala, M. Mutational landscape of cancer-driver genes across human cancers. Sci Rep 13, 12742 (2023). 10.1038/s41598-023-39608-2

27 Canisius, S., Martens, J. W. & Wessels, L. F. A novel independence test for somatic alterations in cancer shows that biology drives mutual exclusivity but chance explains most co-occurrence. Genome Biol 17, 261 (2016). 10.1186/s13059-016-1114-x

28 Wang, X., Kostrzewa, C., Reiner, A., Shen, R. & Begg, C. Adaptation of a mutual exclusivity framework to identify driver mutations within oncogenic pathways. Am J Hum Genet 111, 227–241 (2024). 10.1016/j.ajhg.2023.12.009

29 Iranzo, J., Gruenhagen, G., Calle-Espinosa, J. & Koonin, E. V. Pervasive conditional selection of driver mutations and modular epistasis networks in cancer. Cell Rep 40, 111272 (2022). 10.1016/j.celrep.2022.111272

30 Ciriello, G., Cerami, E., Sander, C. & Schultz, N. Mutual exclusivity analysis identifies oncogenic network modules. Genome Res 22, 398–406 (2012). 10.1101/gr.125567.111

31 Scharpf, R. B. et al. Genomic Landscapes and Hallmarks of Mutant RAS in Human Cancers. Cancer Res 82, 4058–4078 (2022). 10.1158/0008-5472.CAN-22-1731

32 Patterson, A., Elbasir, A., Tian, B. & Auslander, N. Computational Methods Summarizing Mutational Patterns in Cancer: Promise and Limitations for Clinical Applications. Cancers (Basel) 15 (2023). 10.3390/cancers15071958

33 Liu, C., Han, Z., Zhang, Z. K., Nussinov, R. & Cheng, F. A network-based deep learning methodology for stratification of tumor mutations. Bioinformatics 37, 82–88 (2021). 10.1093/bioinformatics/btaa1099

34 Huang, J. K., Jia, T., Carlin, D. E. & Ideker, T. pyNBS: a Python implementation for network-based stratification of tumor mutations. Bioinformatics 34, 2859–2861 (2018). 10.1093/bioinformatics/bty186

35 Hofree, M., Shen, J. P., Carter, H., Gross, A. & Ideker, T. Network-based stratification of tumor mutations. Nat Methods 10, 1108–1115 (2013). 10.1038/nmeth.2651

36 Zhang, W., Ma, J. & Ideker, T. Classifying tumors by supervised network propagation. Bioinformatics 34, i484–i493 (2018). 10.1093/bioinformatics/bty247

37 Killock, D. Genetics: HotNet2-see the wood for the trees. Nat Rev Clin Oncol 12, 66 (2015). 10.1038/nrclinonc.2014.234

38 Reyna, M. A., Leiserson, M. D. M. & Raphael, B. J. Hierarchical HotNet: identifying hierarchies of altered subnetworks. Bioinformatics 34, i972–i980 (2018). 10.1093/bioinformatics/bty613

39 Lin, C.-J. On the convergence of multiplicative update algorithms for nonnegative matrix factorization. IEEE Transactions on Neural Networks 18, 1589–1596 (2007).

40 Wang, Y.-X. & Zhang, Y.-J. Nonnegative matrix factorization: A comprehensive review. IEEE Transactions on knowledge and data engineering 25, 1336–1353 (2012).

41 Bajpai, A. K. et al. Systematic comparison of the protein-protein interaction databases from a user’s perspective. J Biomed Inform 103, 103380 (2020). 10.1016/j.jbi.2020.103380

42 Milano, M., Agapito, G. & Cannataro, M. Challenges and Limitations of Biological Network Analysis. BioTech (Basel) 11 (2022). 10.3390/biotech11030024

43 Alexandrov, L. B. et al. The repertoire of mutational signatures in human cancer. Nature 578, 94–101 (2020). 10.1038/s41586-020-1943-3

44 Consortium, A. P. G. AACR Project GENIE: Powering Precision Medicine through an International Consortium. Cancer Discov 7, 818–831 (2017). 10.1158/2159-8290.CD-17-0151

45. Zhao, W. X., et al. A survey of large language models. arXiv preprint arXiv:2303.18223 1 (2023).

46 Vaswani, A. et al. Attention is all you need. Advances in neural information processing systems 30 (2017).

47 Devlin, J., Chang, M.-W., Lee, K. & Toutanova, K. in Proceedings of the 2019 conference of the North American chapter of the association for computational linguistics: human language technologies, volume 1 (long and short papers). 4171-4186.

48 O’Neil, N. J., Bailey, M. L. & Hieter, P. Synthetic lethality and cancer. Nat Rev Genet 18, 613–623 (2017). 10.1038/nrg.2017.47

49 Levine, J. H. et al. Data-Driven Phenotypic Dissection of AML Reveals Progenitor-like Cells that Correlate with Prognosis. Cell 162, 184–197 (2015). 10.1016/j.cell.2015.05.047

50 Zheng, F. et al. Interpretation of cancer mutations using a multiscale map of protein systems. Science 374, eabf3067 (2021). 10.1126/science.abf3067

51 Kong, J. et al. Prediction of immunotherapy response using mutations to cancer protein assemblies. Sci Adv 10, eado9746 (2024). 10.1126/sciadv.ado9746

52 Hanahan, D. Hallmarks of cancer-Then and now, and beyond. Cell (2026). 10.1016/j.cell.2025.12.049

53 Liberzon, A. et al. The Molecular Signatures Database (MSigDB) hallmark gene set collection. Cell Syst 1, 417–425 (2015). 10.1016/j.cels.2015.12.004

54 Galan-Cobo, A. et al. KEAP1 and STK11/LKB1 alterations enhance vulnerability to ATR inhibition in KRAS mutant non-small cell lung cancer. Cancer Cell 43, 1530–1548 e1539 (2025). 10.1016/j.ccell.2025.06.011

55 Wohlhieter, C. A. et al. Concurrent Mutations in STK11 and KEAP1 Promote Ferroptosis Protection and SCD1 Dependence in Lung Cancer. Cell Rep 33, 108444 (2020). 10.1016/j.celrep.2020.108444

56 Liu, J. et al. Wnt/beta-catenin signalling: function, biological mechanisms, and therapeutic opportunities. Signal Transduct Target Ther 7, 3 (2022). 10.1038/s41392-021-00762-6

57 Thorsson, V. et al. The Immune Landscape of Cancer. Immunity 48, 812–830 e814 (2018). 10.1016/j.immuni.2018.03.023

58 Park, S. et al. A deep learning model of tumor cell architecture elucidates response and resistance to CDK4/6 inhibitors. Nat Cancer 5, 996–1009 (2024). 10.1038/s43018-024-00740-1

59 Papillon-Cavanagh, S., Doshi, P., Dobrin, R., Szustakowski, J. & Walsh, A. M. STK11 and KEAP1 mutations as prognostic biomarkers in an observational real-world lung adenocarcinoma cohort. ESMO Open 5 (2020). 10.1136/esmoopen-2020-000706

60 Knetki-Wroblewska, M., Wojas-Krawczyk, K., Krawczyk, P. & Krzakowski, M. Emerging insights into STK11, KEAP1 and KRAS mutations: implications for immunotherapy in patients with advanced non-small cell lung cancer. Transl Lung Cancer Res 13, 3718–3730 (2024). 10.21037/tlcr-24-552

61 Blattner, M. et al. SPOP Mutation Drives Prostate Tumorigenesis In Vivo through Coordinate Regulation of PI3K/mTOR and AR Signaling. Cancer Cell 31, 436–451 (2017). 10.1016/j.ccell.2017.02.004

62 Swami, U. et al. SPOP Mutations as a Predictive Biomarker for Androgen Receptor Axis-Targeted Therapy in De Novo Metastatic Castration-Sensitive Prostate Cancer. Clin Cancer Res 28, 4917–4925 (2022). 10.1158/1078-0432.CCR-22-2228

63 Jamaspishvili, T. et al. Clinical implications of PTEN loss in prostate cancer. Nat Rev Urol 15, 222–234 (2018). 10.1038/nrurol.2018.9

64 Majumder, A., Meyer, B. S., Hicks, J. K., Boyle, T. A. & Haura, E. B. CTNNB1 mutation can mediate resistance to EGFR, ALK and KRAS targeted therapies. Cancer Research 82, 1101–1101 (2022).

65 Du, Z. et al. Molecular mechanisms of acquired resistance to EGFR tyrosine kinase inhibitors in non-small cell lung cancer. Drug Resist Updat 82, 101266 (2025). 10.1016/j.drup.2025.101266

66 Kulhavy, J. et al. Impact of Baseline beta-Catenin Comutations on Prognosis in EGFR-Mutant Lung Cancer. JCO Precis Oncol 9, e2400771 (2025). 10.1200/PO-24-00771

67 Di Conza, G., Ho, P. C., Cubillos-Ruiz, J. R. & Huang, S. C. Control of immune cell function by the unfolded protein response. Nat Rev Immunol 23, 546–562 (2023). 10.1038/s41577-023-00838-0

68 Ayers, M. et al. IFN-gamma-related mRNA profile predicts clinical response to PD-1 blockade. J Clin Invest 127, 2930–2940 (2017). 10.1172/JCI91190

69 Wu, J. N. & Roberts, C. W. ARID1A mutations in cancer: another epigenetic tumor suppressor? Cancer Discov 3, 35–43 (2013). 10.1158/2159-8290.CD-12-0361

70 Dhar, S. S. & Lee, M. G. Cancer-epigenetic function of the histone methyltransferase KMT2D and therapeutic opportunities for the treatment of KMT2D-deficient tumors. Oncotarget 12, 1296–1308 (2021). 10.18632/oncotarget.27988

71 Okamura, R. et al. ARID1A alterations function as a biomarker for longer progression-free survival after anti-PD-1/PD-L1 immunotherapy. J Immunother Cancer 8 (2020). 10.1136/jitc-2019-000438

72 Li, J. et al. Epigenetic driver mutations in ARID1A shape cancer immune phenotype and immunotherapy. J Clin Invest 130, 2712–2726 (2020). 10.1172/JCI134402

73 Wang, D. X. et al. Mutation status of the KMT2 family associated with immune checkpoint inhibitors (ICIs) therapy and implicating diverse tumor microenvironments. Mol Cancer 23, 15 (2024). 10.1186/s12943-023-01930-8

74 Wang, G. et al. CRISPR-GEMM Pooled Mutagenic Screening Identifies KMT2D as a Major Modulator of Immune Checkpoint Blockade. Cancer Discov 10, 1912–1933 (2020). 10.1158/2159-8290.CD-19-1448

75 Minton, K. Predicting variant pathogenicity with AlphaMissense. Nat Rev Genet 24, 804 (2023). 10.1038/s41576-023-00668-9

76 Torkamani, A., Wineinger, N. E. & Topol, E. J. The personal and clinical utility of polygenic risk scores. Nat Rev Genet 19, 581–590 (2018). 10.1038/s41576-018-0018-x

77 Boehm, K. M., Khosravi, P., Vanguri, R., Gao, J. & Shah, S. P. Harnessing multimodal data integration to advance precision oncology. Nat Rev Cancer 22, 114–126 (2022). 10.1038/s41568-021-00408-3

78 Capper, D. et al. DNA methylation-based classification of central nervous system tumours. Nature 555, 469–474 (2018). 10.1038/nature26000

79 Cunningham, H., Ewart, A., Riggs, L., Huben, R. & Sharkey, L. Sparse autoencoders find highly interpretable features in language models. arXiv preprint arXiv:2309.08600 (2023).

80 Ferber, D. et al. Development and validation of an autonomous artificial intelligence agent for clinical decision-making in oncology. Nat Cancer 6, 1337–1349 (2025). 10.1038/s43018-025-00991-6

81 Truhn, D. et al. Artificial intelligence agents in cancer research and oncology. Nat Rev Cancer (2026). 10.1038/s41568-025-00900-0

82 Xian, S. et al. Transformer patient embedding using electronic health records enables patient stratification and progression analysis. NPJ Digit Med 8, 521 (2025). 10.1038/s41746-025-01872-z

83 Lipkova, J. et al. Artificial intelligence for multimodal data integration in oncology. Cancer Cell 40, 1095–1110 (2022). 10.1016/j.ccell.2022.09.012

84 Ballard, J. L., Wang, Z., Li, W., Shen, L. & Long, Q. Deep learning-based approaches for multi-omics data integration and analysis. BioData Min 17, 38 (2024). 10.1186/s13040-024-00391-z

85 Ye, X., Shi, T., Huang, D. & Sakurai, T. Multi-Omics clustering by integrating clinical features from large language model. Methods 239, 64–71 (2025). 10.1016/j.ymeth.2025.03.017

86 Kundra, R. et al. OncoTree: A Cancer Classification System for Precision Oncology. JCO Clin Cancer Inform 5, 221–230 (2021). 10.1200/CCI.20.00108

87 Choudhury, N. J. et al. The GENIE BPC NSCLC Cohort: A Real-World Repository Integrating Standardized Clinical and Genomic Data for 1,846 Patients with Non-Small Cell Lung Cancer. Clin Cancer Res 29, 3418–3428 (2023). 10.1158/1078-0432.CCR-23-0580

88 Sanz-Garcia, E. et al. Genomic Characterization and Clinical Outcomes of Patients with Peritoneal Metastases from the AACR GENIE Biopharma Collaborative Colorectal Cancer Registry. Cancer Res Commun 4, 475–486 (2024). 10.1158/2767-9764.CRC-23-0409

89 Samstein, R. M. et al. Tumor mutational load predicts survival after immunotherapy across multiple cancer types. Nat Genet 51, 202–206 (2019). 10.1038/s41588-018-0312-8

90 Giordano, T. J. The cancer genome atlas research network: a sight to behold. Endocr Pathol 25, 362–365 (2014). 10.1007/s12022-014-9345-4

91 Van Allen, E. M. et al. Genomic correlates of response to CTLA-4 blockade in metastatic melanoma. Science 350, 207–211 (2015). 10.1126/science.aad0095

92 Snyder, A. et al. Genetic basis for clinical response to CTLA-4 blockade in melanoma. N Engl J Med 371, 2189–2199 (2014). 10.1056/NEJMoa1406498

93 Hugo, W. et al. Genomic and Transcriptomic Features of Response to Anti-PD-1 Therapy in Metastatic Melanoma. Cell 165, 35–44 (2016). 10.1016/j.cell.2016.02.065

94 Miao, D. et al. Genomic correlates of response to immune checkpoint therapies in clear cell renal cell carcinoma. Science 359, 801–806 (2018). 10.1126/science.aan5951

95 Hellmann, M. D. et al. Genomic Features of Response to Combination Immunotherapy in Patients with Advanced Non-Small-Cell Lung Cancer. Cancer Cell 33, 843–852 e844 (2018). 10.1016/j.ccell.2018.03.018

96 Rizvi, N. A. et al. Cancer immunology. Mutational landscape determines sensitivity to PD-1 blockade in non-small cell lung cancer. Science 348, 124–128 (2015). 10.1126/science.aaa1348

97 Zhao, J. et al. Immune and genomic correlates of response to anti-PD-1 immunotherapy in glioblastoma. Nat Med 25, 462–469 (2019). 10.1038/s41591-019-0349-y

98 Miao, D. et al. Genomic correlates of response to immune checkpoint blockade in microsatellite-stable solid tumors. Nat Genet 50, 1271–1281 (2018). 10.1038/s41588-018-0200-2

99 Chu, D. & Wei, L. Nonsynonymous, synonymous and nonsense mutations in human cancer-related genes undergo stronger purifying selections than expectation. BMC Cancer 19, 359 (2019). 10.1186/s12885-019-5572-x

100 Chakravarty, D. et al. OncoKB: A Precision Oncology Knowledge Base. JCO Precis Oncol 2017 (2017). 10.1200/PO.17.00011

101 Lin, Z. et al. Evolutionary-scale prediction of atomic-level protein structure with a language model. Science 379, 1123–1130 (2023). 10.1126/science.ade2574

102 Le, D.-H. in 2017 4th NAFOSTED Conference on Information and Computer Science. 242–247 (IEEE).

103 Lee, D. et al. Pan-cancer mutational signature analysis of 111,711 targeted sequenced tumors using SATS. medRxiv (2024). 10.1101/2023.05.18.23290188

104 Rosales, R. A., Drummond, R. D., Valieris, R., Dias-Neto, E. & da Silva, I. T. signeR: an empirical Bayesian approach to mutational signature discovery. Bioinformatics 33, 8–16 (2017). 10.1093/bioinformatics/btw572

105 Bagaev, A. et al. Conserved pan-cancer microenvironment subtypes predict response to immunotherapy. Cancer Cell 39, 845–865 e847 (2021). 10.1016/j.ccell.2021.04.014

